# Cancer tissue of origin constrains the growth and metabolism of metastases

**DOI:** 10.1101/2022.08.17.504141

**Authors:** Sharanya Sivanand, Yetis Gultekin, Peter S. Winter, Sidney Y. Vermeulen, Konstantine Tchourine, Keene L. Abbott, Laura V. Danai, Florian Gourgue, Brian T. Do, Kayla Crowder, Tenzin Kunchok, Allison N. Lau, Alicia M. Darnell, Alexandria Jefferson, Satoru Morita, Dan G. Duda, Andrew Aguirre, Brian M. Wolpin, Nicole Henning, Virginia Spanoudaki, Laura Maiorino, Darrell J. Irvine, Omer H. Yilmaz, Caroline A. Lewis, Dennis Vitkup, Alex K. Shalek, Matthew G. Vander Heiden

## Abstract

Metastases arise from a subset of cancer cells that disseminate from the primary tumor; however, the factors that contribute to proliferation of cancer cells in a secondary site are incompletely understood. The ability of cancer cells to thrive in a new tissue site is influenced by genetic and epigenetic changes that are important for disease initiation and progression, but these factors alone do not predict if and where cancers metastasize. Specific cancer types metastasize to consistent subsets of tissues, suggesting that factors associated with the primary tumor influence the tissue environments where cancers can grow. Using pancreatic cancer as a model, we find that primary and metastatic tumors share many metabolic similarities and that the tumor initiating capacity and proliferation of both primary- and metastasis-derived cells is favored in the primary site relative to the metastatic site. Moreover, propagating lung or liver metastatic cells in vivo to enrich for tumor cells adapted to grow in the lung or the liver does not enhance their relative ability to form large tumors in those sites, change their preference to grow in the primary site, nor stably alter some aspects of their metabolism relative to primary tumors. We also analyzed primary liver and lung cancer cells and find that these cells also exhibit a preference to grow in their primary site relative to metastatic sites. Together, these data suggest that the cancer tissue-of-origin influences the metabolism of both primary and metastatic tumors and may impact whether cancer cells can thrive in a metastatic site.

## Introduction

Metastasis contributes to the high mortality of patients with cancer. Metastases arise from a subset of cancer cells within the primary tumor^1^; however, why some clones thrive in new tissue sites and what determines which tissue sites will support proliferation of metastatic cancer cells is incompletely understood. It is known that formation of metastasis is a rare event^2^. The fact that metastases are derived from a subset of cells from the primary tumor may suggest that only those cancer cells that are adapted to grow in the new tissue site are selected for during metastasis. Oncogenic mutations are important contributors to primary tumor initiation and disease progression^3^, but causal genetic determinants of metastasis have not been identified. This has led to the speculation that epigenetic alterations must be involved in allowing cancer cells to thrive in new tissue sites; however, consistent gene expression programs that predict which cancers can grow in any specific tissue have also not been found. These data argue that genetic factors alone do not predict metastasis^4^, and the fact that cancers arising in different tissue sites tend to metastasize to distinct subsets of tissue locations that are characteristic of that tumor type argues that some property of the primary tumor influences the specific tissues to where cancer cells can metastasize. Nevertheless, the properties of the primary tumor environment that are shared with the metastatic tumor environment to allow metastasis are not well understood.

One factor that could be shared between primary and metastatic sites and might influence whether tumors can grow in each location is nutrient availability, as metabolism is a property of cancer that is influenced by tissue environment^5–7^. Genetic events, such as oncogenic *Ras* signaling, contribute to metabolic changes in cancer^8–10^; however, cancer tissue-of-origin and tumor location also influence cancer metabolic phenotypes^11,12^. The nutrients available to cancer cells in tumors depends on tissue location and cancer type^13–15^, and tumor cells can exhibit some metabolic plasticity to adapt aspects of metabolism to proliferate in a metastatic site^16,17^. Indeed there is evidence that specific metabolic adaptations are required to grow in some tissues^18–20^; however, tumor metabolic gene expression better resembles the tissue-of-origin for a cancer than it does tumors arising in other tissues^21,22^, and differences in nutrient availability across tissues might also constrain the tissue of origin-shaped metabolism of cancer cells and limit where cells can thrive as metastases following tumor colonization^13,21–23^. This model would predict some metabolic similarities are retained between the primary and metastatic tumors, and that some aspects of the metabolism of the metastatic tumor is influenced by the tissue-of-origin, a possibility that has not been tested. This led us to investigate how well primary- and metastasis-derived cancer cells grow in different tissues, and how metabolism of primary and metastatic tumors relates to the preferences to grow in different tissue sites.

### Metastases are derived from a subset of cells

Pancreatic ductal adenocarcinoma (PDAC) has a high incidence of metastasis, and the genetically engineered *LSL-Kras^G12D/+^; Trp53^R^*^172^*^H/+^; Pdx-1-Cre* (KPC) mouse pancreatic cancer model recapitulates many features of the human disease including a propensity to metastasize^24,25^. Over a period of 6-8 months, KPC mice develop tumors in the pancreas and frequently develop metastatic lesions in the liver, and occasionally in the lung^24^, a metastatic pattern similar to that observed in patients. To study primary and metastatic PDAC cells, cancer cells were isolated from primary tumors and matched liver or lung metastases that arose in the KPC model, and that proliferated at similar rates when cultured in standard conditions in vitro (Supplemental Fig. 1a-c). We also verified that the cells express mutant Kras (Supplemental Fig. 1d). To confirm past observations that cells derived from metastases have an increased ability to colonize the organ to which it had metastasized^20,26^, cells derived from matched PDAC primary tumors or liver metastases were engineered to express either mCherry or GFP, such that equal numbers of cells expressing different fluorescent proteins could be mixed and implanted into the pancreas, liver, or flank (subcutaneous) in syngeneic C57BL/6J mice (Supplemental Fig. 2a). We confirmed that the labeled cell populations had approximately equivalent representation prior to implantation (Supplemental Fig. 2b), and that a fixed number of cells could form tumors of similar size in the pancreas when injected individually, or co-injected as a mixed population, despite expressing different fluorescent proteins (Supplemental Fig. 2c). When a mixed population of cells derived from primary tumors and liver metastases were implanted in the pancreas, the resulting tumors were enriched for primary tumor-derived cells when analyzed by either flow-cytometry or immunohistochemistry, and this observation held regardless of the fluorophore expressed (Supplemental Fig. 2d) even though mCherry expression is known to be more immunogenic than GFP expression^27^. As expected, when a mixed population of cells was implanted in the liver, liver metastases-derived cells were more abundant regardless of whether they expressed mCherry or GFP, even though primary tumor-derived cells also contributed to the resulting tumor (Supplemental Fig. 2g-i). Interestingly, when a mixed population of primary- and liver metastasis-derived cells were implanted in the flank, the resulting tumor was derived primarily from one of the two cell populations, and this was not determined by which fluorescent protein was expressed (Supplemental Fig. 2j, k). These data are consistent with tumors at either the primary or a metastatic site being derived from a subset of cancer cells, and are consistent with numerous studies showing that cancer cells derived from a specific metastatic site have an increased ability to metastasize to that same site^28^. They also confirm that the cancer cells lines generated exhibit known phenotypes associated with cancer cells derived from metastases.

That implantation of primary and metastatic cancer cells into the flank, a site not previously experienced by either primary or liver-metastatic PDAC cells, results in expansion of tumors derived predominantly from a subset of cells (Supplemental Fig. 2j, k) suggests that stochastic processes, perhaps related to metastasis being a rare event,^2^ may contribute to which cells contribute to the tumor^1,29,30^. To test this possibility further, we labeled primary pancreatic cancer cells with either mCherry or GFP, mixed them in equal proportions (Fig. 1a), and implanted the mixed population into the pancreas, liver, or flank. Even though the different labeled cancer cells were derived from the same cell populations, in most cases either an mCherry or GFP labeled cell was found to be the dominant contributor to tumors that formed regardless of site, and which labeled cell predominated in the resulting tumor was not affected by which fluorescent protein was expressed (Fig. 1b-d). These data are consistent with observations of clonal dominance in pancreatic cancer in both primary and metastatic sites^31^, but also raise the possibility that the presence of subclones from the primary tumor in metastases reflects, at least in part, that subsets of cells contribute to a bulk tumor in different locations.

**Figure 1.**
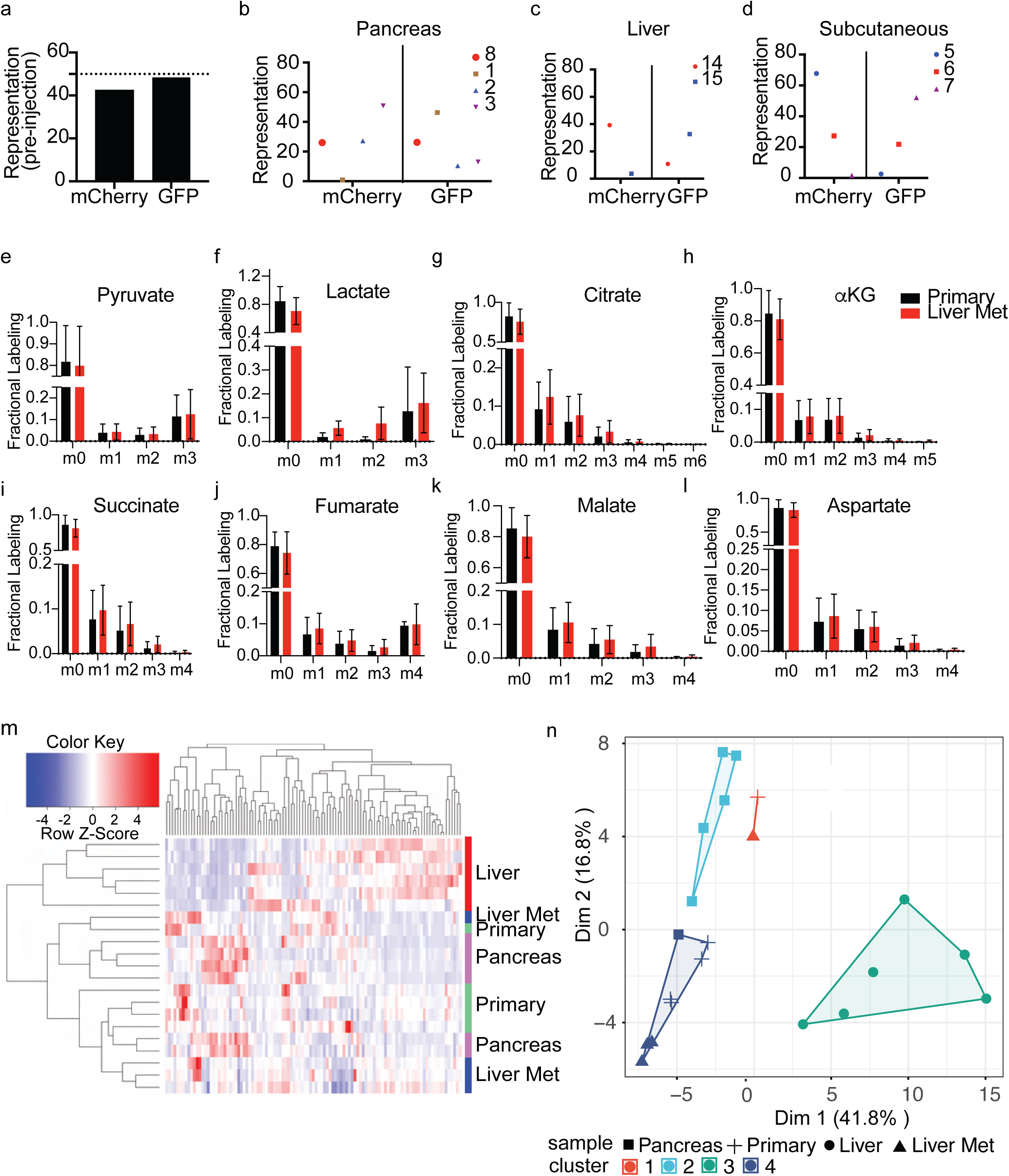
Primary and metastatic pancreatic tumors exhibit some similar metabolic phenotypes. (**a)** The same cancer cells isolated from a primary pancreatic tumor arising in a KPC mouse were engineered to express either mCherry or GFP and combined in equal numbers prior to implantation into different tissue sites in mice. Flow cytometry confirming that approximately equal representation of each labeled cancer cell population is present in the mixed population. (**b-d**) Representation of mCherry and GFP labeled cells in tumors derived from injection of the mixed population shown in (a) into the pancreas (n=4) (b), liver (n=2) (c), or subcutaneous space (n=3) (d). (**e-l**) Paired pancreatic cancer cells derived from primary tumors or liver metastases arising in the KPC mouse model were implanted into the pancreas or liver, respectively, and the resulting tumor-bearing mice were infused for 6 hours with U-^13^C glucose at 0.4 mg/min to assess glucose fate in tumors growing in each site. Fractional labeling of the indicated metabolites as determined by GC-MS is shown. Data are from 3-4 mice per group. Mean +/- stdev. (**m-n**) Relative metabolite levels in autochthonous paired primary pancreatic cancer and liver metastases arising in KPC mice were assessed by LC-MS; Tumors were harvested in the mornings at the same time of the day. Metabolite levels were also measured for normal pancreas and liver tissue from age-matched non-tumor bearing control mice. The metabolite data for each sample is clustered in two different ways: Unsupervised clustering represented as a heatmap (m) or K-means unsupervised clustering (n); 6 mice were used for the normal tissue analysis and 4 mice were used for the paired primary tumor and metastasis analysis.

### Glucose and glutamine contribution to metabolism is similar in primary and metastatic tumors

Several studies have noted that the availability of specific nutrients can determine whether cancer cells can thrive in a metastatic site^32^; however, those studies have focused on how metabolism of a specific tissue metastasis might differ from a primary tumor, and a comprehensive analysis of both the metabolic similarities and differences for primary and metastatic tumors growing in different sites is lacking. To begin to assess whether accessing a metastatic tissue environment with sufficient similarity to the primary site is important to support the metabolism of metastatic cancer cells, we implanted primary- or liver metastasis-derived PDAC cells into the pancreas or liver to form tumors in mice. Assessment of transplanted tumors was necessary to study the metabolism of metastases, because it was not possible to isolate large enough metastatic tumors from the KPC model for functional analysis of metabolism. Analysis of implanted tumors also allows comparison of tumors within a predictable time window that enables analysis of isotopically-labeled glucose fate in tumor tissue in conscious, unrestrained tumor bearing mice^33^. Mice were analyzed after a 6 hour ^13^C-glucose infusion, a time where metabolite labeling approaches steady state allowing comparison of glucose fate between tissues^33^. Similar ^13^C-glucose enrichment was observed in plasma of labeled-glucose infused mice bearing either pancreatic tumors or liver metastatic tumors (Supplemental Fig. 3a), and minimal differences in glucose fate were observed when comparing how metabolites were labeled in tumors growing in the pancreas or liver (Fig. 1e-l).

To extend these findings to a second mouse model of pancreatic cancer, we utilized cancer cells isolated from the *LSL-Kras^G12D/+^; Trp53^-/-^; Pdx-1-Cre* (KP^-/-^C) mouse PDAC model^34,35^, and implanted those cells into the pancreas, liver, or flank to again assess the fate of U-^13^C glucose in tumors arising in each location, as well as glucose fate in the normal pancreas and liver of mice without tumors. While a different labeling pattern was observed when comparing metabolites from tumors to metabolites from the normal pancreas or liver, minimal differences in metabolite labeling was observed in pancreatic tumors growing in different tissue sites (Supplemental Fig. 3b-i).

We also assessed the fate of infused U-^13^C glutamine in tumors arising from KP^-/-^C cells that were implanted in either in the pancreas or the liver and again found minimal differences in metabolite labeling in tumors growing in these two sites (Supplemental Fig. 4a-f). Together, these data argue that in the pancreatic cancer models studied, there are many similarities in how glucose and glutamine are metabolized when comparing primary and metastatic tumors even though glucose metabolism in the tumors differs from that observed in normal pancreas and normal liver.

### Metabolite levels are similar in primary and metastatic pancreatic tumors

To further study how tissue site influences the metabolism of pancreatic cancer liver metastases, we compared metabolite levels in primary pancreatic tumors, matched liver metastases, normal pancreas, and normal liver. For these studies, we assessed metabolite levels in the tissues harvested from age-matched normal as well as tumor bearing mice. Unsupervised clustering of metabolites suggest that the metabolic profile of normal liver is most distinct from the other samples (Fig. 1m). Moreover, this type of analyses revealed less separation between primary tumors, liver metastases, and the normal pancreas relative to the normal liver when the same data was analyzed using two different approaches (Fig. 1m, n). Of note, although we observe variability across samples, we do not find evidence of consistent changes in metabolites that distinguish primary and liver metastatic tumors from KPC mice (Supplemental Fig. 5a). Taken together with the data assessing glucose and glutamine fate, these data suggest that pancreatic tumors retain some similarities in metabolic phenotype that are shared between the primary site and liver metastasis, and that the metabolic phenotype of a liver metastatic pancreatic tumor more closely resembles the primary tumor and normal tissue-of-origin than it does the metastatic tissue.

We next examined relative metabolite levels in cultured pancreatic cancer cells derived from primary tumors and liver metastases to determine whether the similarities are recapitulated in purified cancer cells isolated from each site. We find metabolite levels were similar across different independently derived paired primary and metastatic cells such that clustering based on metabolites did not uniformly segregate primary cells from liver metastatic cells (Supplemental Fig. 6a). To examine whether nutrient utilization differs between primary- and liver metastasis-derived cancer cells in culture, we examined glucose metabolism by assessing the fate of U-^13^C-glucose. Although some differences in glucose fate in culture were observed across select cells derived from primary and liver metastases, these differences were not consistent across multiple paired primary and liver metastatic lines from independent mice (Supplemental Fig 6b-j). Taken together, these data argue that any major metabolic differences that exist between primary and liver metastatic PDAC cells are not maintained in standard culture, and thus are not determined by stable genetic or epigenetic control of metabolism, although these data should not be interpreted to say there are no metabolic differences between primary pancreatic tumors and liver metastases.

### Cancer cells from primary and metastatic tumors grow best in the primary site

If the need to access a tissue environment with enough similarity to the primary tissue site is a barrier to metastasis, we reasoned that this may result in metastatic cancer cells retaining a preference to grow in the primary site. For instance, if a specific nutrient environment is needed to support the growth of tumors in either the primary or metastatic sites, this environment would be better represented in the primary tissue and result in differences in the rate of tumor growth in different sites. Prior studies of metastasis have focused on how tumor metabolism impacts whether cancer cells can exit the primary tumor, survive in circulation, or find and grow in a particular site^32,36^. However, to our knowledge, the preferential ability of cancer cells derived from primary and metastatic tumors to expand in different environments upon colonization has been largely unexplored. To examine this, we implanted cells derived from either primary tumors or matched liver metastases from KPC mice into the pancreas or liver, as well as into the flank as a neutral site that is commonly used to assess tumor growth in mice (Fig. 2a). In all cases, the cells lacked any fluorophore expression, and the same number of cells were implanted in each site prior to analysis of tumor size after a fixed period of four weeks. We assessed tumor weight where possible, or if too small to accurately weigh, we assessed the weight of the tumor bearing organ as well as the tissue weight of the corresponding normal tissue from age-matched mice. We found that cancer cells from both primary and liver metastatic tumors were able to form tumors at all sites but observed a clear preference for cancer cells derived from both primary tumors and liver metastasis to form tumors in the pancreas, with much larger tumors forming in this organ than in the liver or flank (Fig. 2b, Supplemental Fig. 7a-b). Interestingly, cancer cells formed tumors that grew to a similar size in each organ site regardless of whether the cells were derived from a primary or a liver-metastatic lesion. Tumors in each site were also histologically similar regardless of whether they were derived from a primary tumor or a liver metastasis and histological tumor grade was also similar to naturally arising primary and metastatic tumors in KPC mice (Supplemental Fig. 7c,d,f). Cell proliferation as determined by either Ki67 or BrdU staining was comparable within tumors formed from primary- or liver metastasis-derived cells in both the pancreas and liver (Fig. 2c,e, Supplemental Fig. 8a,d). We also assessed cleaved caspase 3 staining as a marker of cell death in the same tumors, and again found comparable staining within tumors formed from primary- or liver metastasis-derived cells in both the pancreas or liver (Fig. 2f, Supplemental Fig. 8e-f). Of note, regardless of whether the tumor was derived from primary- or liver metastasis-derived cells, both Ki67 and BrdU staining trended lower, and cleaved caspase 3 staining trended higher in tumors growing in the liver compared to tumors growing in the pancreas, a finding consistent with larger tumors forming in the pancreas. Repeating the transplantation experiments with an independently derived matched tdTomato-expressing primary and liver metastasis cell pair, where tdTomato tumor cells from both primary tumors and liver metastasis were derived from mice where a *LSL-tdTomato* reporter allele was bred to the KPC PDAC model, also showed a preference for both primary- and liver metastasis-derived cells to form large tumors in the pancreas (Supplemental Fig. 9a). These data argue that the pancreas better supports the growth of pancreatic cancer cells as tumors, even if those cancer cells are derived from liver metastasis.

**Figure 2.**
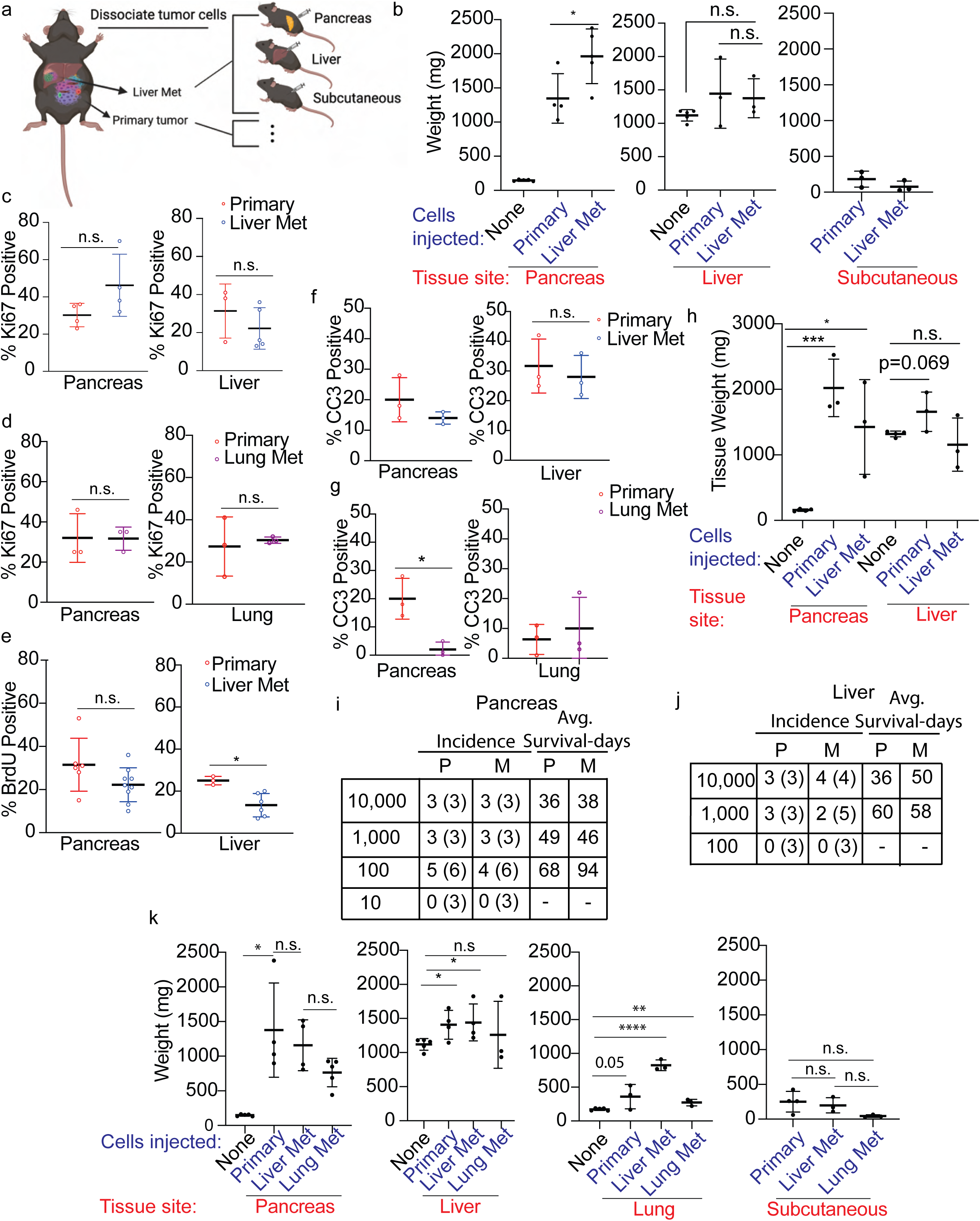
Pancreatic cancer cells derived from both the primary tumor and from metastatic sites generate tumors that grow fastest in the pancreas. (**a**) Schematic depicting transplantation experiments to quantitatively assess pancreatic cancer cell proliferation in different tissue sites. (**b**) Equal numbers of cancer cells isolated from primary pancreatic tumors or paired liver metastases arising in the KPC mouse model were implanted into the pancreas, liver, or subcutaneous space, and resulting tumor size assessed after four weeks. Tissue site indicates the site where cells were implanted, and cells injected indicates whether the cells injected were derived from a primary tumor or liver metastasis (liver met). Relative weights of tumor and associated normal tissue compared with normal tissue of age-matched mice, or tumor weight in the subcutaneous flank, is shown. n=3-5 per group. Male mice were used for all the comparisons. Mean+/- stdev is shown. * p < 0.05; n.s. – not significant. (**c-d**) The percent of cells that stain positive for Ki67 by immunohistochemistry was quantified in tissue sections from tumors arising from primary pancreatic cancer or liver metastatic cells implanted into the pancreas or liver as indicated (c) or implanted into the pancreas or lung as indicated (d). Data from 4-5 mice is shown; Ki67 percentage = number of positive stained cells in each image field divided by the total number of cells in the same field x 100. One representative field per tumor was analyzed; n.s.- not significant; mean +/- stdev is shown (**e**) The percent of cells stained positive for BrdU as assessed by immunohistochemistry was quantified in tissue sections from tumors arising from primary or metastatic cells in either the pancreas or the liver as indicated; each data point represents one region of a tumor obtained from n=2-3 mice per group; Quantification was determined by manually counting number of positive cells divided by the total number of cells in the same field x 100; * p <0.05; mean +/- stdev is shown (**f-g**) The percent of cells that stain positive for cleaved caspase 3 by immunohistochemistry was quantified on tissue sections derived from tumors arising from either primary or metastatic cells in either the pancreas or liver (f) or in tumors arising from primary or metastatic cells in either the pancreas or the lung (g). Each data point indicates an average of three regions from each mouse (n=3 mice/group). Percent positivity was quantified by manually counting number of positive cells divided by the total number of cells in the same field x 100. n.s.- not significant; * p <0.05 mean +/- stdev (**h**) Relative weights of tumors derived from equal numbers of cancer cells implanted into nude (NU/J) mice after four weeks and associated normal tissue weights from age-matched mice is shown. As in (b), tissue site indicates the site where cells were implanted, and cells injected indicates whether the cells injected were derived from a primary tumor or liver metastasis. n=3-4 mice/group; both male and female mice were used. Mean +/- stdev is shown. *p<0.05, ***p<0.001, n.s.- not significant. (**i-j**) The indicated number of cancer cells derived from a primary pancreatic tumor (P) or liver metastases (M) were implanted into either the pancreas (i) or the liver (j), and animals followed to determine if a tumor formed as well as how long mice with tumors survived post-implantation; male mice were used for all the conditions. The number of mice with tumors is shown, with the total number of mice injected for each number of cells indicated in parentheses. These data were also used to calculate an approximate tumor initiating capacity (TIC) for primary- and metastasis-derived cells at each tissue site: Pancreas, primary (1/63.4), liver met (1/99.2), p-value = 0.531; Liver, primary (1/417), liver met (1/2005), p-value= 0.106. (**k**) Equal numbers of cancer cells isolated from primary pancreatic tumors or paired liver or lung metastases arising in the KPC mouse model were implanted into the pancreas, liver, lung (via tail vein), or subcutaneous space, and resulting tumor size assessed after four weeks. As in (b), relative weights of tumor and associated normal tissue compared with normal tissue of age-matched mice, or tumor weight in the subcutaneous flank, is shown, and tissue site indicates the site where cells were implanted, and cells injected indicates whether the cells injected were derived from a primary tumor, liver, or lung metastasis. n=3-5 per group, Mean+/- stdev is shown. * p < 0.05, ** p< 0.05, and **** p < 0.0001; n.s. – not significant.

### A preference to grow in the primary site is independent of a T cell mediated immune response

The fact that immunogenic tdTomato fluorophore expression did not affect the preference for both primary- and liver metastasis-derived cells to grow in the pancreas argues against a tissue-specific difference in adaptive immune response explaining this phenotype; however, to test this possibility further we implanted the same number of primary tumor or liver-metastasis derived PDAC cells into the pancreas or liver to form tumors over a fixed time in an immunocompromised host. Paired primary and metastatic cells were injected into nude mice that lack T cells, and again we observed that both primary tumor- and liver metastasis-derived PDAC cells formed larger tumors in the pancreas (Fig. 2h, Supplemental Fig. 9b). Weight loss was noted in some nude mice following tumor implantation that was not observed when cells were implanted into syngeneic hosts. To control for this, we normalized tumor size data to body weight, and still observe a preference for both primary tumor- and liver metastasis-derived cells to grow in the pancreas compared to the liver in nude mice (Supplemental Fig. 9c-d). These data argue that a preference for PDAC cells grow in the pancreas relative to the liver cannot be explained by tissue specific differences in a T cell-mediated immune response.

### Fewer cells are required to form tumors in the primary site

To examine whether the tumor initiating capacity of primary- and liver metastasis-derived cells differ in different tissue locations, we implanted different numbers of either primary- or liver metastasis-derived pancreatic cancer cells into the pancreas or liver and determined the minimal number of cells required to form tumors in each site. We found that both primary- and liver metastasis-derived cells have a similar tumor initiating capacity at each site; however, more cells are needed to form a tumor in the liver than are needed to form tumors in the pancreas regardless of whether they are derived from a primary or a metastatic pancreatic tumor (Fig. 2i,j). These data further support the notion that pancreatic cancer cells retain a preference to grow in the primary site even if they are derived from a liver metastasis.

### Pancreatic cancer cells retain a preference to grow in the primary site regardless of the site of metastasis

To assess whether PDAC lung metastatic cells also retain a preference to grow in the pancreas, we performed similar experiments using independently derived cells that lack any fluorophore expression from matched primary, liver, and lung metastases arising in KPC mice (Supplemental Fig. 7e). Cells derived from either liver metastases or lung metastases can form tumors in both the liver and lung; however, by far the largest tumors developed in the pancreas with the smallest tumors in the flank (Fig. 2k, Supplemental Fig. 10a-f). The tumors forming in the lung were also histologically similar regardless of whether they were derived from a primary tumor or a metastasis (Supplemental Figure 7f). Further, tumors formed from primary- or lung metastasis-derived cells showed similar staining for markers of proliferation and cell death when growing in the lung or the pancreas (Fig. 2d,g and Supplemental Fig. 8b,c,e,g). These data argue pancreatic cancer cells from metastasis arising in the lung also retain a preference to grow in the primary site.

Propagation of cancer cells in metastatic sites has been used to select for cancer cells to grow in a particular tissue site to study metastasis, including assessment of metabolic differences between primary and metastatic tumors^37–39^. Of note, these studies have largely examined the ability of selected cells to seed metastases when implanted in the primary site^39^, and it remains unclear whether cells selected to seed metastatic sites also improve in their ability to grow in the different tissue sites once they arrive at that site. To answer this latter question, tumor cells derived from lung or liver metastases arising in KPC mice were propagated as tumors in mice by three rounds of repeated implantation and passaging in the lung or liver respectively (Fig. 3a, Supplemental Fig. 11a). Consistent with published results^37,38^, lung metastatic cancer cells selected for in this manner efficiently formed lung tumors when injected via tail vein (Supplemental Fig. 11b), and natural lung metastases developed in mice when these cells were implanted in the pancreas to form tumors in the primary site (Supplemental Fig. 11c). These same properties were also present in the parental cells derived from a natural lung metastasis (Supplemental Fig. 11b, c), which are otherwise infrequent in the KPC model^24^. Of note, spontaneous lung metastases were never observed from primary- or liver metastasis-derived cells implanted into either the pancreas or the liver. Propagating liver metastatic cancer cells as liver tumors in vivo resulted in more efficient tumor formation when the cells were implanted in the liver or when injected via tail vein (Supplemental Fig. 11d, e). Of note, in a defined time frame, formation of liver nodules was not observed following tail vein injection of parental cells derived from liver metastases or following tail vein injection of primary- or lung metastasis-derived cells, confirming that in vivo passaging of cancer cells in the liver results in an increased ability to seed new liver tumors as reported in other contexts^37^.

**Figure 3.**
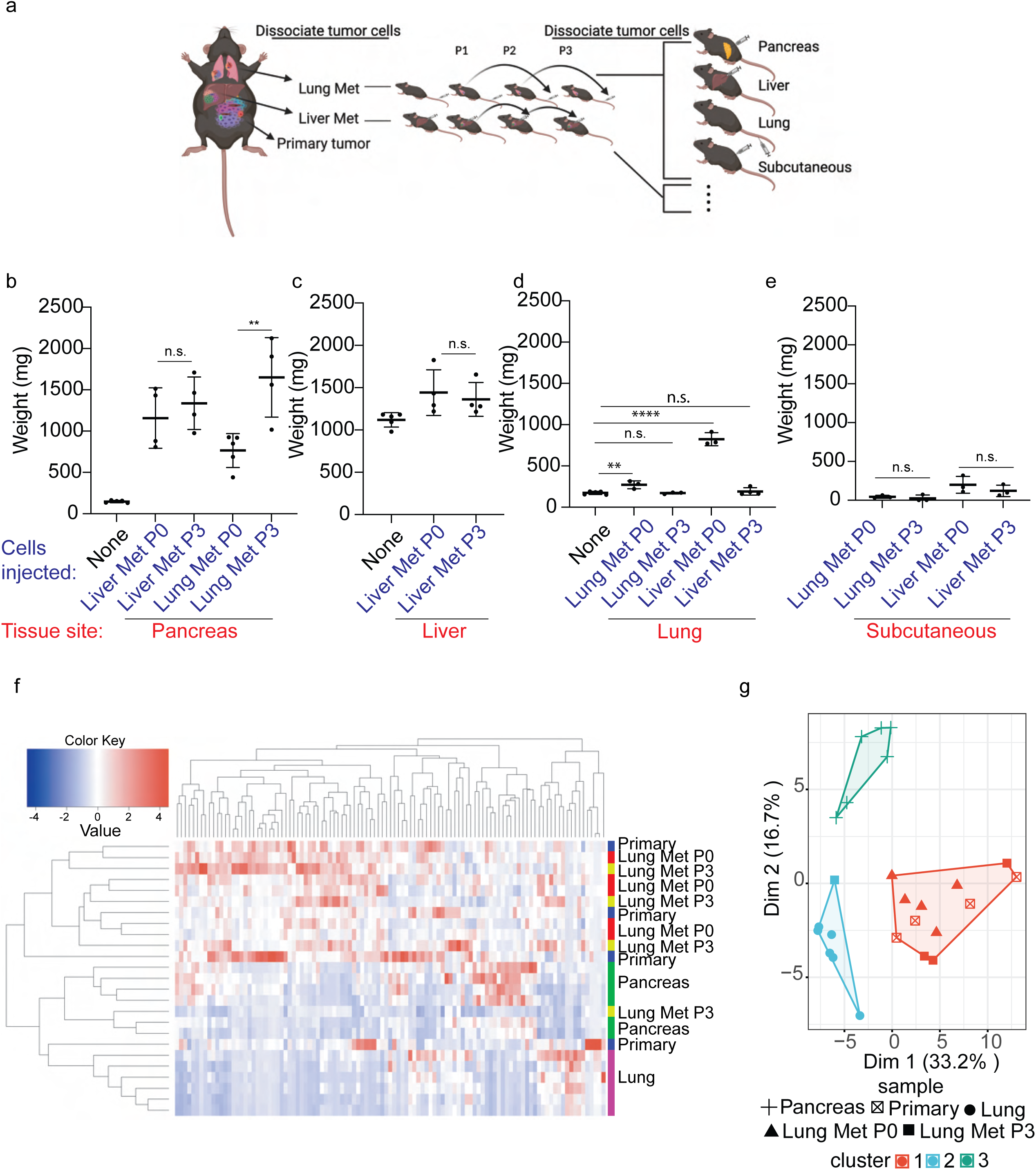
Pancreatic cancer cells retain metabolic phenotypes found in the primary tumor even when repeatedly passaged in a metastatic site. (**a**) Schematic depicting propagation of lung and liver pancreatic cancer metastases arising in KPC mice by implantation into the lung or liver to form tumors three times prior to use in transplantation experiments to quantitatively assess proliferation and metabolic phenotypes in different tissue sites. (**b-e**) Equal numbers of cancer cells isolated from primary pancreatic tumors, paired liver, or lung metastases (P0), or liver or lung metastatic cancer cells that were passaged in the liver or lung as described in (a) (P3) were implanted into the pancreas, liver, lung (via tail vein), or subcutaneous space, and resulting tumor size assessed after 3 weeks. Relative weights of tumor and associated normal tissue compared with normal tissue of age-matched mice, or tumor weight in the subcutaneous flank, is shown. Tissue site indicates the site where cells were implanted, and cells injected indicates whether the cells injected were derived from a P0 or P3 liver or lung metastasis. Mean+/- stdev; n.s.- not significant, ** p<0.005; *p <0.05; n=3-5 mice per group. (**f-g**) Relative metabolite levels arising in tumors from primary, P0 and P3 lung metastases cells implanted in the pancreas (primary) or lung were assessed by LC-MS. Metabolite levels were also measured for normal pancreas and liver tissue from age-matched wild-type mice. The metabolite data for each sample is clustered in two different ways: Unsupervised clustering represented as a heatmap (f) or K-means unsupervised clustering (g).

Despite passaging cancer cells as tumors in the lung or liver, when their ability to form tumors in different tissue sites was quantitatively assessed over a defined time window, the liver- and lung-selected cells still formed larger tumors in the pancreas (Fig. 3b-e, Supplemental Fig. 11f). Again, cells derived from both liver and lung metastases were able to form tumors at all sites, with the largest tumors forming in the pancreas and the smallest in the flank. Notably, similar size tumors were observed at each site regardless of where the cells were derived from, and whether they were passaged previously as tumors in the liver or lung. These data support a model where pancreatic cancer cells retain a preference to grow in the primary site, even when repeatedly passaged in a metastatic tissue site.

We next examined whether propagation of tumor cells in different tissues alters their metabolic phenotype. We first examined metabolites extracted from tumors that were generated from cells derived from spontaneous lung metastases (P0) and from cells derived from in vivo selected lung metastases (P3) and compared those to metabolite levels extracted from primary pancreatic tumors as well as from age-matched normal lung or normal pancreas tissue. The small size of tumors that formed in the lung prevented assessment of metabolite levels when tumors are growing in that site, however we could assess metabolites extracted from the large tumors that formed in the pancreas to determine whether propagating cells in the lung stably selects for tumors with major alterations in metabolism as these cells retain an ability to spontaneously reseed lung metastases (Supplemental Fig. 11c). We find that unsupervised clustering representation of the data as a heat map revealed that the pancreatic tumors derived from tumors that were generated from lung metastases, spontaneous or selected, cluster based on metabolite levels together with the primary tumor (Fig. 3f). When the same dataset was clustered using unsupervised K-means clustering, one tumor derived from passaging of cancer cells in the lung clustered with the normal lung while the remaining tumor samples clustered together (Fig. 3g). These data suggest that passaging cancer cells as tumors in the lung, for the most part, does not select for extensive stable alterations in metabolism that are retained when these cells are grown in the pancreas, although should not be interpreted to mean that there are no differences in metabolites present between the tumors and the normal tissues.

### A preference to grow in the primary tissue site is observed for both primary lung and primary liver cancers

To assess whether a preference for cancer cells to form tumors in the pancreas might be explained by the pancreas microenvironment being more permissive to tumor growth, we asked whether cancer cells derived from different cancers also have a preference to grow in the pancreas relative to their primary tissue site. First, we considered primary lung adenocarcinoma cells derived from the *LSL-Kras^G12D/+^; Trp53^fl/fl^; Ad-Cre* mouse lung cancer model^40^. Primary lung cancer cells did not grow well as tumors in the pancreas, yet these cells formed large tumors in the lungs of syngeneic mice (Fig. 4a). Of note, this preference to grow in the lung relative to the pancreas was the opposite of what we observed with pancreatic cancer lung metastasis-derived cells (Fig. 3b,d and Supplemental Fig. 12a). We next considered whether direct delivery of cancer cells into the lung provides the same results as initiating lung tumors via tail vein injection. First, we delivered the same number of primary lung cancer cells or PDAC lung metastasis-derived cells via intratracheal injection and observed after a fixed time that primary lung cancer-derived cells form more nodules than pancreatic lung cancer metastasis-derived cells (Fig. 4b, Supplemental Fig. 12b). Next, we directly injected the same number of primary lung cancer cells or primary PDAC cancer cells directly into the lung parenchyma and observed that after a fixed time the primary lung cancer-derived cells give rise to larger tumors than primary pancreatic cancer cells (Supplemental Fig. Fig. 12c,d).

**Figure 4.**
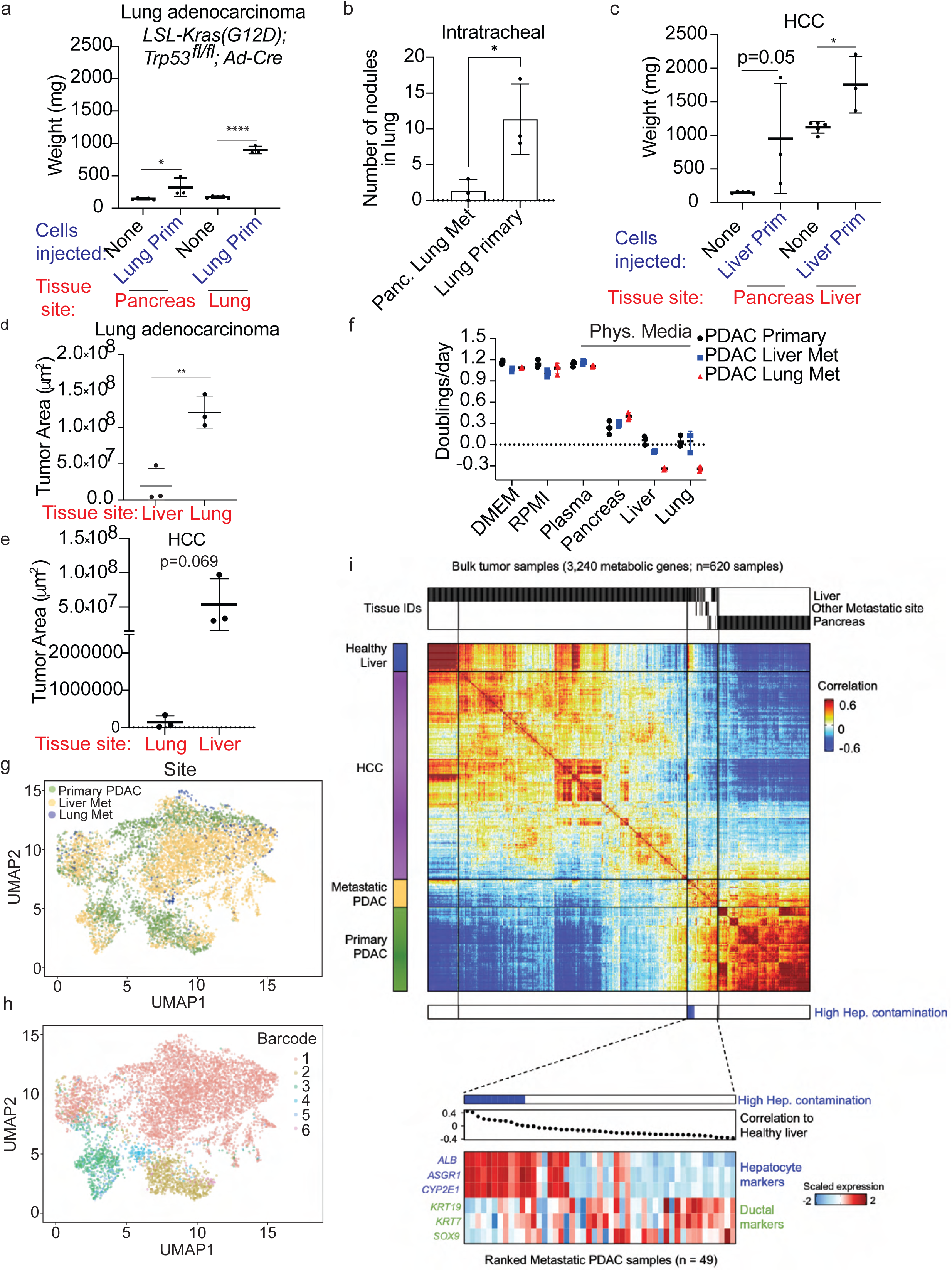
Tissue-of-origin influences the metabolism and tissue site preference for metastatic tumor growth. (**a**) Equal numbers of lung adenocarcinoma cells derived from the *LSL-Kras^G12D/+^; Trp53^fl/fl^; Ad-Cre* mouse model were implanted into the pancreas or delivered to the lung and resulting tumor size assessed after four weeks. Relative weights of tumor and associated normal tissue compared with normal tissue from age-matched mice is shown. Mean+/- stdev; * p<0.05, **** p<0.0001; n=3-6 mice per group. (**b**) Equal number of cells derived from pancreatic lung metastasis (Lung Met) or from primary lung cancer described in (a) were introduced into the lung via intratracheal inhalation of the cancer cells; tumor burden in the lung is depicted as number of lung nodules in each section of 3 mice/group. Each data point represents one mouse. Mean +/- stdev; * p <0.05. (**c**) Equal numbers of RIL-176 hepatocellular carcinoma (HCC) cells were implanted into the pancreas or the liver and resulting tumor size assessed after four weeks. Relative weights of tumor and associated normal tissue compared with normal tissue from age-matched mice is shown. Mean+/- stdev; * p<0.05. (**d**) Equal numbers of primary lung cancer cells described in (a) were implanted into the liver or delivered to the lung via tail vein injection and tumor area determined via analysis of histological sections after 4 weeks. Each data point represents one mouse. Mean +/- stdev. **p <0.005 (**e**) Equal numbers of primary HCC cells described in (c) were implanted into the liver or delivered to the lung via tail vein injection and tumor area in each site was quantified as in (d) after 4 weeks. (**f**) Proliferation rate of primary pancreatic, liver, or lung metastatic cells in either standard cell culture media (DMEM, RPMI) or in media formulated to match the metabolite concentrations measured in normal mouse plasma or in mouse tissue (pancreas, liver, or lung) interstitial fluid. Representative data from at least n=2 independent experiments per line per condition is shown. mean +/- stdev; * p <0.05, ** p <0.001, **** p < 0.0001. (**g-h**) UMAP plot comparing metabolic gene expression in single cells from primary PDAC, lung metastases (lung met), and liver metastases (liver met) from the KPC model (g); and the UMAP plot of metabolic gene expression analysis in (g) with each clonal populations represented; these clones are arbitrarily labeled as clones 1-6 and are do not match clone numbers in the original publication (h). (**i**) Cross correlation of bulk metabolic gene expression profiles involving 3,240 genes from 620 samples with gene expression obtained from either hepatocellular carcinoma (HCC) TCGA, PDAC (Panc-seq), or PDAC TCGA data (top). Ranked metastatic samples with high degree of hepatocyte contamination as assessed by higher gene expression of hepatocyte markers and lower gene expression of ductal markers is shown on the bottom with corresponding correlation to normal liver.

To assess whether hepatocellular carcinoma (HCC)-derived cells have a preference to grow in the liver relative to the pancreas, a fixed number of mouse HCC cells^41,42^ were implanted into the pancreas or liver of syngeneic mice, and tumor burden assessed after four weeks. While there was some variability between mice, HCC cells implanted into liver trended toward larger tumors relative to tumors that formed in the pancreas even after accounting for normal tissue weight (Fig. 4c, Supplemental Fig. 12e). However, the same number of HCC-derived cells formed larger tumors over a fixed period of time in the liver than pancreatic cancer liver metastasis-derived cells (Fig. 3b,c and Supplemental Fig. 12e). The pancreas is not a typical site of metastasis for HCC or primary lung cancer although in patients these cancers can metastasize to the lung and liver, respectively. Thus, we next asked whether primary HCC and primary lung cancer cells retain a preference to grow in their respective primary tissue sites relative to the lung or liver. When a fixed number of primary lung cancer cells are transplanted into the liver or delivered to the lung, we observe that after a fixed time the lung cancer cells form larger tumors in the lung than they do in the liver (Fig. 4d; Supplemental Fig. 13a). Similarly, the HCC cells form larger tumors in the liver than they do in the lung (Fig. 4e; Supplemental Fig. 13b). These data support a model where cancer cells from different origins retain a preference to grow in their tissue of origin relative to a metastatic site.

### Primary and metastatic cancer cells proliferate better in nutrient conditions that match those found in the primary site

To begin to test whether differences in tissue nutrient availability contribute to the preference of cancer cells to grow in their primary site, we first measured the absolute concentrations of metabolites in interstitial fluid isolated from normal mouse tissues as well as matched plasma^13^. The concentration of most metabolites in tissue interstitial fluid were lower than those measured in plasma, and what nutrients were present in interstitial fluid differed depending on tissue site (Supplemental Fig. 14a). Next, we formulated media to match the nutrient levels measured in plasma or interstitial fluid from each tissue site and assessed the ability of cells to proliferate in these media relative to standard DMEM- or RPMI-based culture media. We find that primary pancreatic cancer cells, pancreatic cancer liver metastasis-derived cells, and pancreatic cancer lung metastasis-derived cells proliferate at similar rates in standard media or media with plasma nutrient condition (Fig. 4f). While proliferation was much slower in media formulated to match nutrient levels measured in tissues, the pancreatic cancer primary- and metastasis-derived cells proliferated at similar rates in media with pancreas nutrient levels but fail to expand in the time frame assayed in media with lung or liver nutrient conditions. While we were unable to expand any cells in media with liver interstitial fluid nutrient levels, media with lung interstitial nutrient levels can support proliferation of some cancer cells. Both primary mouse lung cancer cells as well as established A549 human lung cancer cells proliferate in lung interstitial fluid nutrient conditions in contrast to pancreatic cancer lung metastasis-derived cells (Supplemental Fig. 14b). Taken together, these data support a model where nutrient levels in the primary tissue can better support the proliferation of cancer cells that arise in that tissue than nutrient levels found in metastatic sites.

### Primary and metastatic tumors express similar metabolic genes

To further examine whether metastatic tumors retain some aspects of metabolism as tumors growing in the primary site, we queried an available dataset to determine whether metabolic gene expression is conserved between primary tumors, liver metastases, and lung metastases that arise in the KPC mouse pancreatic cancer model^31^. Analysis of metabolic genes from this single-cell RNA-seq (scRNA-seq) dataset revealed significant overlap in metabolic gene expression between primary PDAC and liver metastases when all cells were analyzed (Supplemental Fig. 15a). Of note, when the analysis was restricted to the most abundant clonal populations found in the tumor samples in the dataset, despite evidence for heterogeneity in metabolic gene expression among cancer cells isolated from both the primary tumor and the metastases, similar cell heterogeneity in metabolic gene expression appeared to be present in cells derived from both the primary and metastatic sites, although very few cells derived from lung metastases were available for analysis (Fig. 4g,h, Supplemental Fig. 15b). Interestingly, when analyzed via the barcodes in the dataset that allow the cancer cells injected that were implanted in the pancreas to be traced across tissues^31^, cells that cluster by barcode and do not segregate by tissue site when considering metabolic gene expression (Fig. 4g,h, Supplemental Fig. 15c). Thus, metabolic heterogeneity in the pancreatic cancer cells from this dataset is driven more by clonal relationships between cells than by the tissue environment where the cells were found.

That is, these data argue that despite heterogeneity in metabolic gene expression among pancreatic cancer cells isolated from each tissue site, and metabolic heterogeneity among cancer cell clones, we did not find evidence for selection of a clone defined by a specific global metabolic gene expression pattern in cells derived from metastases. Rather, these data suggest that PDAC cancer cells arising in KPC mice retain a similar global metabolic gene expression program to support tumor growth regardless of site, although they do not argue against specific metabolic adaptations being associated with tumor growth in a particular site.

To assess whether these same findings related to metabolic gene expression are found in human cancers, we analyzed expression of metabolic genes from available pancreatic cancer and HCC patient tumor-derived RNA-seq datasets^43,44^. Consistent with tissue-of-origin having a stronger influence on global metabolic gene expression than tissue site, we find significant overlap of metabolic gene expression between primary pancreatic tumors and liver metastatic pancreatic tumors, and that this metabolic gene expression is distinct from that observed in primary liver cancer, which overlaps with normal liver tissue (Fig. 4i, Supplemental Fig. 15d-e). Notably, the human pancreatic cancer liver metastases that overlap most with primary liver cancer samples contain a higher degree of contamination with normal hepatocytes as determined by higher expression of hepatocyte markers (Fig. 4i, Supplemental Fig. 15d-e). These data further support a model where metastatic cancer cells retain many aspects of the metabolic program found in the primary tumor, and that this may constrain cancer cell proliferation in metastatic tissue sites.

## Discussion

The finding that many aspects of the metabolic program found in metastases are shared with those found in the primary tumor argues that the metabolic program cancer cells use to support proliferation is in part defined by the tissue-of-origin even when exposed to a new metastatic tissue environment. Of note, these data do not rule out that some phenotypes are selected for to enable cancer cells to proliferate in metastatic sites. There is evidence that specific metabolic adaptions can promote growth of cancer in specific tissues^15,45–49^; however it appears that the metabolic plasticity of cancer cells may not be as flexible as often assumed^17^. Rather, that cancers prefer to grow in the primary site supports a model where the inflexible portions of a metabolic program derived from the cancer tissue-of-origin might limit the nutrient environments where cancer cells can grow. A relative lack of metabolic plasticity may explain why chemotherapies that target metabolism remain effective in treating both primary and metastatic tumors, with patients selected for treatment based on the cancer tissue-of-origin. Moreover, this model may underlie, at least in part, why particular cancer types metastasize to stereotyped locations, as accessing a nutrient environment that is similar enough to the primary tumors may be necessary to support some parts of the metabolic program that is selected for within the primary tumor. Nevertheless, there is limited evidence from a small number of patients that some metastases from primary renal cell cancer, breast cancer, and colorectal cancer exhibit faster radiographic progression than primary tumors or have increased Ki67 staining compared to the primary tumor^50,51^. Radiographic progression of individual lesions in patients is often observed in patients with later stage cancers and could be influenced by multiple factors including prior therapies the patients received. Metastases to contralateral unaffected healthy tissue that matches the primary site is observed in some cancers, such as lung cancer, but is not common in other cancers such as breast cancer. Whether some cancer types exhibit more extensive metabolic adaptation upon metastases than the pancreatic, lung, and hepatocellular cancers considered here remains unknown. Regardless, better understanding the impact of different tissue nutrient environments on proliferation of cancers arising in different sites could inform the selection of treatment modalities for patients based on the pattern of metastasis for a given primary tumor.

## Materials and Methods

### Animal studies

All studies were approved by the MIT committee on Animal Care under protocol number #0119-001-22. For autochthonous models, *Kras^G12D/+^; Trp53^R^*^172^*^H/+^; Pdx1-Cre* (KPC)^24^ and *Kras^G12D/+^; Trp53^R^*^172^*^H/+^; Pdx1-Cre; LSL-tdTomato* (KPCT)^33^ mice from a mixed 129/Sv and C57Bl6/J background as well as pure C57Bl6/J genetic background were used for most experiments. For the experiments involving immunocompromised mice, both male and female 8-10 week old nude mice (NU/J) obtained from Jackson labs were used. Both male and female pure C57Bl6/J mice were used for all other transplantation experiments. For metabolomics experiments normal tissue was isolated from age-matched 6-month-old mice, while tumors and paired liver metastases were isolated from the same 6–8-month-old animals. Tissues for metabolite measurements were isolated at the same time of day. Animals were housed under 12-hour light and 12-hour dark cycle, and co-housed with littermates with ad libitum access to water and food, except immediately following surgical procedures.

### Tumor transplantation

Both male and female animals were used for these studies. For subcutaneous, pancreatic, or intrahepatic transplantation studies, C57B16/J or Nu/J mice aged approximately 6-8 weeks (for C57B16/J) or 8-10 weeks (for Nu/J) were injected with 100,000 cells PDAC cells derived from primary or metastatic tumors arising in KPC or KPCT mice at the indicated site.

Cells were delivered into the pancreas using established protocols as previously published^33^. To introduce cells into the liver, briefly, upon anesthetization, a small incision was introduced to exteriorize one lobe of the liver and cells were directly injected into the liver. Mice were monitored for recovery and appropriate post-operative care was provided for all surgical procedures in line with our approved animal protocols. To deliver cancer cells to the lung via circulation, 100,000 cells were injected into the tail vein. Mice were euthanized 4 weeks post injection of tumor cells or at signs of animal distress. All mice within the same experimental group were euthanized at the same time point. For direct ultrasound-guided lung tumor induction to deliver cells into the lung parenchyma, 10,000 cells were injected into 18 week old male C57BL6/J mice. In brief, mice are first depilated and imaged by microCT (Skyscan 1276 x-ray microtomograph (Bruker)) in pronated position to get three axis coordinates for injection in the left lung using distances from three landmarks: liver-lung interface, pleural line, and the skin. Animals are then transferred to the Vevo3100 ultrasound imager (FujiFilm-VisualSonics), keeping the mice in pronated position. Identification of the three landmarks by ultrasound imaging combined with the use of syringe mounted on a precision driver allows for delivery of 2 µl of cell suspension leading to development of a focal nodule in the left lung. Mice were euthanized three weeks post injection of tumor cells.

For intratracheal delivery of cancer cells into lungs, a variation of a previously published protocol^40^ was used. Briefly, anesthetized mice were placed on a specialized platform suspended by their front teeth such that the chest was vertical. A light source was used for visualization and a catheter (Clint; CP-26746) was guided into the open trachea. Following placement of the catheter, 50 μl cell suspension in HBSS was pipetted into the lung (total cell number per mouse delivered was 100,000) with inhalation of the cell suspension noted based on observed mouse inhalation and disappearance of the liquid in the catheter.

### Limiting dilution studies to assess tumor initiation

For limiting dilution studies to assess tumor initiation cells, mice were injected with the indicated number of cells and monitored twice a week for signs of tumor burden. The tumor initiating capacity was calculated using ELDA software (http://bioinf.wehi.edu.au/software/elda/). At least 3 mice were included for each condition.

### Cell competition experiments

For cell competition experiments, pancreatic cancer cells derived from primary or liver metastatic tumors were engineered to express either mCherry or GFP and mixed in equal numbers. Pre-injection representation was confirmed using flow cytometry (BD LSR-II). 100,000 cells containing the mixed population were injected into the pancreas, liver, or subcutaneously, and after tumors formed, they were excised, digested, and analyzed by flow cytometry to determine relative representation in the tumor. Alternatively, tumor tissue was obtained, fixed for histology and relative representation of mCherry or GFP expressing cells assessed by immunohistochemistry.

### Tumor adaptation to grow in the lung and liver

For metastatic site adaptation experiments, cells from naturally arising liver and lung metastatic tumors in the KPC mouse model were isolated and cultured for less than 10 population doublings. Cells from the lung metastases were then transplanted into the lung via tail vein injection. Resulting tumors were dissociated and re-transplanted into secondary recipient mice without *in vitro* propagation. This process was repeated three times. A similar approach was taken for liver metastases where cells were implanted in the liver, with resulting tumors dissociated and re-transplanted in secondary recipient mice without *in vitro* propagation three times. Mice were euthanized at different time points based on established and approved criteria, and following the last round of *in vivo* selection, tumors were dissociated and cultured for less than 5 population doublings before use in transplantation experiments (P0 refers to the parental cells (prior to *in vivo* adaptation); P3 refers to the tumor and tumor cells derived from three round of *in vivo* adaptation) to grow in the liver or lung as indicated.

### Cell isolation and cell culture

Cells were isolated from primary and metastatic mouse pancreatic tumors as described previously^33^. Briefly, tumors were exteriorized, minced, and digested with collagenase XI (Sigma C9407) and dispase II (Roche 04942078001) and plated in DMEM. RIL-175 mouse hepatocellular carcinoma cells (HCC) were isolated from the Duda Laboratory at MGH and derived from hepatic tumors established in C57BL/6 mice as previously described^44^. Lung adenocarcinoma cells were obtained from the *LSL-*Kras(G12D); *Trp53^fl/fl^; Ad-Cre* mouse lung cancer model as previously described^52^. Cells were cultured in DMEM (Corning 10-013-CV) supplemented with 10% heat inactivated fetal bovine serum. Penicillin-streptomycin was added only at the time of cell isolation from mice. Cells were regularly tested for mycoplasma contamination using the MycoAlert Plus kit (Lonza).

### Cell proliferation

30,000 cells were plated in a 6 well plate in 2 mL of DMEM with 10% FBS and cultured for at least 12 h. Cells were washed once, and media was replaced with fresh DMEM at the time of cell counting on day 0 using a Cellometer Auto T4 Plus Cell Counter (Nexcelom Bioscience). Cells counted again 3 days later, and doublings per day was calculated using the formula: Proliferation rate (Doublings/day) = Log_2_(Final cell count (day 3)/initial cell count (day 0))/3 (days).

### Isotope labeling experiments in cultured cells

Cells were plated in 6-well plates, and the next day cells were washed three times with warm PBS, and DMEM without glucose and pyruvate supplemented with 10% dialyzed FBS and 10 mM U-^13^C-glucose (Cambridge Isotope Laboratories) was added for 24 hours prior to metabolite extraction.

### Isotope labeled nutrient infusion experiments in mice

Infusion of U-^13^C glucose or U-^13^C glutamine (Cambridge Isotope Laboratories) into control or tumor bearing mice was performed as previously described^33,53^. Three-weeks after implantation of cancer cells, animals underwent surgical catheter implantation in the jugular vein 3-4 days prior to labeled glucose infusion. Mice were fasted for 4 hours prior to starting the infusion and animals remained conscious and mobile for the duration of the infusion. Labeled glucose was delivered at a rate of 0.4 mg/min for 6 hours, and then plasma and tumor tissue were isolated and flash frozen for analysis by mass spectrometry. Labeled glutamine was delivered at a rate of 3.7 mg/kg/min for 6 hours. All isotope labeling experiments in mice were performed at the same time of day.

### BrdU incorporation in tumors

BrdU (10 mg/mL in PBS) was injected intraperitoneally (I.P.) into mice in 200 μL volume per 20 g of mouse body weight. 24 hours post injection, tumors and associated normal tissue was collected and fixed in 10% formalin prior to staining with anti-BrdU antibody (Abcam; ab6326) using immunohistochemistry at a dilution of 1:2000 in PBS.

### Metabolite Extraction

To analyze glucose in plasma, 10 μL of plasma was extracted with 100% methanol and dried down with nitrogen and derivatized with 50 μL of 2 wt% hydroxylamine hydrochloride (2% Hox) in pyridine followed by incubation at 90°C for 60 min. 100 μL of propionic anhydride was added, and samples were incubated at 60°C for 30 min, followed by evaporation under nitrogen at room temperature overnight. The next day, dried samples were dissolved in 100 μL of ethyl acetate and transferred to glass vials for analysis by GC-MS.

For tissue metabolite analysis, the harvested tissues were rinsed briefly in ice-cold blood blank saline and flash frozen in liquid nitrogen. Frozen tissues were then ground into powder using a pre-chilled mortar and pestle. The tissue powder was then weighed into pre-chilled tubes and extracted with methanol (containing 500 nM each of 17 isotopically labeled ^13^C/^15^N amino acids (Cambridge Isotope Laboratories, Inc.)): chloroform: water (6:3:4 v/v/v), vortexed for 10 min and centrifuged for 10 min at maximum speed. Polar metabolites were transferred to Eppendorf tubes, dried under nitrogen gas, and resuspended in different volumes of water containing labeled non-standard amino acid mix (Cambridge Isotope Laboratories, MSK-NCAA-1) to account for differences in starting tissue weight.

For cultured cells, cells were seeded at 30,000 cells/well in a 6 well dish in 2 mL of medium and incubated for 72 hour, or 100,000 cells were plated and incubated overnight. Media was aspirated from cells and then washed rapidly in ice cold bank saline followed by addition of 500 mL ice-cold 80% methanol in water containing 500 nM each of 17 isotopically labeled ^13^C/^15^N amino acids (Cambridge Isotope Laboratories). Alternatively, for metabolite analysis by GC-MS, cells were extracted in equal parts 80% methanol (containing 2.5 ng/mL norvaline internal standard) and chloroform. Samples were vortexed 10 min at 4°C and spun at 16,000 xg for 10 min at 4°C. Equal volume of the polar fraction was transferred to a new tube and dried down under nitrogen and frozen at -80°C prior to analysis by mass spectrometry.

### Isolation of tissue interstitial fluid

Interstitial fluid was collected from mouse tissues using an adapted protocol based on prior published work^13^. Plasma was also isolated from the same mice. Male mice on ad libitum diet were used for all tissue collection and interstitial fluid isolation. Organs from 5 pooled mice were combined per isolation which was done was in three different groups for a total of 15 mice. Each pooled sample was treated as an individual data point for the LCMS measurements and analysis. In each group, the age of the mice ranged from 6-8 (cohort 1), 8 weeks (cohort 2), 8 weeks (cohort 3). All mice were euthanized at the same time of day to account for any effects of time of day on metabolism. Tissues were kept on ice throughout the harvest and when ready to pool, were briefly rinsed in ice-cold saline and excess liquid was removed before tissues were placed in a 50 mL conical vial lined with a 20 μm nylon mesh filter (Spectrum Labs, Waltham MA, 148134). The samples were spun at 400 x g for 10 min. The flow through was collected and spun again at 400 x g prior to storage in -80°C until further analysis. Measurements of metabolite levels and absolute concentrations was determined following previously published protocol^13^.

### Formulation of physiological media

To generate media reflecting the measured concentrations of metabolites in tissue interstitial fluid and plasma, individual metabolites were weighed out and combined into different pools; pre-ground pools were further combined to generate the base media. The complete media was comprised of base media supplemented with 10% dialyzed serum. Any metabolite that was not measured in our analysis was substituted with prior measured concentrations found in mouse plasma and was kept consistent across medias.

### Gas chromatography-mass spectrometry

GC-MS was used to analyze metabolites as previously described^33^. Briefly, dried metabolite extracts were dissolved in 16 mL methoxamine (MOX) reagent (ThermoFisher TS-45950) and incubated at 37°C for 90 min. 20 mL N–methyl–N–(tert– butyldimethylsilyl)trifluor-oacetamide + 1% tert–Butyldimethylchlorosilane (Sigma 375934) was added to the sample and incubated at 60°C for 1 hour. Samples were centrifuged for 5 min and 20 mL of the derivatized sample was transferred to GC vial for analysis using a DB-35MS column (Agilent Technologies 122-3832) installed in an Agilent 7890 gas chromatograph coupled to an Agilent 5975C mass spectrometer. The helium carrier gas was used at a constant flow rate of 1.2mL/min. One microliter of the sample was injected at 270°C. After injection, the GC oven was held at 100°C for 1 min, increased to 300°C at 3.5°C/min. The oven was then ramped up to 320°C at 20°C/min and held for 5 min at 320°C. The MS system operated under electron impact ionization at 70 eV and the MS source and quadrupole was held at 230°C and 150°C, respectively. The detector was used in scanning mode and the scanned ion range was 100-650 m/z. Total ion counts were determined by integrating appropriate ion fragments for each metabolite using El-Maven software (Elucidata).

### Liquid chromatography-mass spectrometry

Metabolite profiling was conducted on a QExactive bench top orbitrap mass spectrometer equipped with an Ion Max source and a HESI II probe, which was coupled to a Dionex UltiMate 3000 HPLC system (Thermo Fisher Scientific, San Jose, CA). External mass calibration was performed using the standard calibration mixture every 7 days. Typically, samples were reconstituted in 50 uL water and 2 uL were injected onto a SeQuant® ZIC®-pHILIC 150 x 2.1 mm analytical column equipped with a 2.1 x 20 mm guard column (both 5 mm particle size; EMD Millipore). Buffer A was 20 mM ammonium carbonate, 0.1% ammonium hydroxide; Buffer B was acetonitrile. The column oven and autosampler tray were held at 25^∘^C and 4^∘^C, respectively. The chromatographic gradient was run at a flow rate of 0.150 mL/min as follows: 0-20 min: linear gradient from 80-20% B; 20-20.5 min: linear gradient form 20-80% B; 20.5-28 min: hold at 80% B. The mass spectrometer was operated in full-scan, polarity-switching mode, with the spray voltage set to 3.0 kV, the heated capillary held at 275^∘^C, and the HESI probe held at 350^∘^C. The sheath gas flow was set to 40 units, the auxiliary gas flow was set to 15 units, and the sweep gas flow was set to 1 unit. MS data acquisition was performed in a range of *m/z* = 70–1000, with the resolution set at 70,000, the AGC target at 1x10^6^, and the maximum injection time at 20 msec. Relative quantitation of polar metabolites was performed with TraceFinder™ 4.1 (Thermo Fisher Scientific) using a 5-ppm mass tolerance and referencing an in-house library of chemical standards. Data were filtered according to predetermined QC metrics: CV of pools <25%; R of linear dilution series <0.975. Data was normalized to cell number from a separately plated set of samples collected at the time of metabolite extraction.

For untargeted metabolomics, data were acquired as described above, with ddMS^2^ data collected on pooled samples using a Top-10 method, with stepped collision energies of 15, 30 and 45 V. The resolution was set at 17,500, the AGC target was 21x10^5^, the max IT was 100 ms, and the isolation window was set at 1.0 m/z. Data were analyzed using Compound Discoverer 3.1 (Thermo Fisher Scientific) and by including an in-house mass-list. P-values were adjusted according to the Benjamini-Hochberg method.

### Flow cytometry

Tumors were dissected, minced, and digested at 37°C for 30 min with 1 mg/mL Collagenase I (Worthington Biochemical, LS004194), 3 mg/mL Dispase II (Roche, 04942078001), and 0.1 mg/mL DNase I (Sigma, D4527) in 5 mL PBS. After 30 min, cells were incubated with EDTA to 10 mM at room temperature for 5 min. Cells were filtered through a 70 mm cell strainer, washed twice in PBS, and cells resuspended in flow cytometry staining buffer (Thermo Fischer, 00-4222-57) for fluorescent protein expression analysis on a BD-LSR II flow cytometer. At minimum of 10,000 cells were analyzed per sample.

### Immunohistochemistry

Tissue was fixed in formalin for at least 24 h. Sections from formalin fixed paraffin embedded tissues were stained using antibodies against mCherry (1:500 dilution; Novus Biologicals: NBP1-96752), GFP (1:250 dilution; Novus Biologicals: NB600-308), pan-cytokeratin (1:500 dilution; Abcam: ab133496), Ki67 (1:250 dilution; Novus Biologicals: NB110-89717), BrdU (1:2000 dilution; Abcam: ab6326), or Cleaved Caspase 3 (Asp 175) (1:400 dilution; Cell Signaling 9661). Antibodies were diluted in 10% normal goat serum diluted 1:2 (Thermo Fischer: 50062Z) in PBS-T.

### Histology and image analysis

Histology sections were scanned using Aperio Digital Scanning and imported into web based Aperio eSlide Manager or by importing scanned images into QuPath software. Image analysis was done using Fiji software or using QuPath software. Tumor area and total area was calculated for all the sections and the net tumor area was calculated by dividing the tumor area by the total area. The number of nodules in the lung was calculated manually. Tumor histology grade was assessed using clinical criteria by a gastrointestinal pathologist (O.H.Y.).

### RNA-sequencing analysis of mouse tumors

Metabolic gene expression analysis of mouse tumors was done using pancreatic tumors single cell RNA-sequencing data from a published source^31^. PCA plots depict analysis of metabolic genes from entire cell populations. For mouse single-cell RNA-seq data analysis, after normalizing to total counts, metabolic genes were analyzed from primary, liver, or lung metastases and UMAP plots for the most abundant samples were generated using scanpy version 1.8.0 to determine overlap between gene expression across tissue sites. For single cell analysis, lineage tracing of the most represented cell clones from the study permitted analysis of individual clonal populations and UMAP plots depict overlap between gene expression across overrepresented clones within the tumor across different tissue sites. Pearson correlation was also found between pairs of single cells using the scores for the top 20 PCs.

### RNA-sequencing analysis of human tumors

FASTQs for bulk RNA expression profiles were downloaded from the relevant repository (TCGA, https://toil.xenahubs.net<u>; Metastatic PDAC,</u> dbGaP Study Accession phs001652.v1.p1) and all data were processed using the same pipeline. Briefly, each sample’s sequences were marked for duplicates and then mapped to hg38 using STAR. After running QC checks using RNAseqQC, gene-level count matrices were generated using RSEM. Instructions to run the pipeline are given in the Broad CCLE github repository https://github.com/broadinstitute/ccle_processing. Length-normalized values (TPM) were then transformed according to log2(TPM+1) for downstream analysis. We then scaled and centered the entire dataset to allow relative comparisons across sample types (Normal Liver, HCC, PDAC, and metastatic PDAC).

We tested whether metastatic PDAC more closely resembles the metabolic state of the primary site (pancreas) or the dominant metastatic tissue of residence (liver) and used normal liver and hepatocellular carcinoma (HCC) profiles as relevant comparators. To do this, we trimmed the expression data to 3,240 metabolically associated genes using a literature-curated list. We then performed a cross-correlational analysis (Pearson’s *r*) across all samples and separated this matrix by study after clustering (Ward’s method).

We then generated similarity scores for each metastatic sample (n = 49) by computing their average Pearson’s *r* to either primary PDAC, HCC, or normal liver samples.

### Statistical Analysis

Results are represented as mean +/- standard deviation unless otherwise specified. Statistical analysis was performed using GraphPad Prism Software (Version 9.1.1). Where multiple comparisons were appropriate, 2way ANOVA statistical test was used and statistical significance between multiple comparisons was determined using Tukey’s multiple comparisons test available as part of GraphPad Prism Software Analysis tools. The statistical significance between two groups were calculated using unpaired two-tailed student *t* test where noted.

## Acknowledgements

We would like to thank the MIT Division of Comparative Medicine (DCM) staff and Kylee Pait for help with colony maintenance and animal care, the Koch Institute’s Robert A. Swanson (1969) Biotechnology Center for technical support, specifically the Hope Babette Tang Histology Core, the Flow Cytometry Core, and the Preclinical Imaging and Testing Core Facility. This work was supported in part by the Koch Institute Support (core) Grant P30-CA014051 from the National Cancer Institute. We would also like to thank all members of the Vander Heiden lab for helpful discussions and feedback and note that some schematic images in the figures were created using BioRender.com.

## Author contributions

S.S.: conceptualization, investigation, visualization, methodology, writing-original draft, writing review and editing. Y.G., P.S.W., S.Y.V., K.L.A., A. J., F.G., K.M.T., L.V.D., B.T.D., K.C., T.K., A.N.L., A.M.D., C.A.L.: methodology, investigation, writing-review and editing. S.M., D.G.D.; L.M., N.H., V.S., D.I., O.H.Y: methodology. D.V., B.M.W., A.K.S.: supervision, M.G.V.H.: conceptualization, supervision, visualization, funding acquisition, project administration, methodology, writing-original draft, writing-review and editing. We have included all authors who have contributed to this work.

