## Supplemental Figure Legends for "Cancer tissue of origin constrains the growth and metabolism of metastases"

**Supplemental Figure 1. Proliferation of primary and metastatic pancreatic cancer cells in culture. (a-c)** Proliferation rate (doublings/day) of cells in culture that were isolated from primary tumors, or from liver or lung metastases, as indicated. Data shown were obtained from three independently derived paired cell lines; mean +/- stdev, n=3-4 replicates per cell line. (**d**) Representative western blot analysis assessing mutant Kras expression in protein lysates obtained from the indicated pancreatic cell lines and established human cell line controls and pancreatic stellate cells (PSCs).

**Supplemental Figure 2. Cell competition experiments to form tumors in mice.** (**a**) Schematic of experimental procedure where primary or liver metastatic pancreatic cancer cells (liver met) labeled with mCherry or GFP were injected into the pancreas, liver, or subcutaneous flank; either implanted individually or as a 50:50 mixture of the indicated cells. Representation of each cell population in the final tumor was analyzed by flow cytometry or by immunohistochemistry (IHC). (**b**) Flow cytometry of individually labeled cell populations (top) and a mixed population (bottom left). Approximately equal representation of each labeled cancer cell population in a mixed population was confirmed prior to injection (representative data shown bottom right). (**c**) Tumor weight post injection of individually labeled cancer cells (n=3) or a mixed cell population containing equal numbers of both primary (prim) and liver metastasis (met)-derived pancreatic cancer cells (n=8). Mean +/- stdev. (**d**) Representation of GFP-labeled primary or mCherry-labeled liver metastatic cells in a tumor derived from injection of a mixed population containing equal numbers of labeled primary and liver metastasis-derived pancreatic cancer cells into the pancreas as determined by flow cytometry. Each data point represents one mouse; the numbers associated with each mouse indicate animal ID with each animal being injected with the same mixed population of cells. (**e**) Representation of mCherry-labeled primary or GFP-labeled liver metastatic cells in a tumor derived from injection of a mixed population containing equal numbers of labeled primary and liver metastasis-derived pancreatic cancer cells into the pancreas as determined by flow cytometry. Each data point represents one mouse; the numbers associated with each mouse indicate animal ID with each animal being injected with the same mixed population of cells. (**f**) Immunohistochemistry to assess GFP and mCherry expression in a tumor derived from injection of a mixed population containing equal numbers of primary and liver metastasis-derived pancreatic cancer cells into the pancreas as in (d), although the mixed population of cells were from independently derived primary and liver metastatic cells. Scale bar, 250 μm. (**g**) Representation of GFP-labeled primary or mCherry-labeled liver metastatic cells in a tumor derived from injection of a mixed population containing equal numbers of primary and metastasis-derived pancreatic cancer cells into the liver as determined by flow cytometry. Each data point represents one mouse; the numbers associated with each mouse indicate animal ID with each animal being injected with the same mixed population of cells. (**h**) Immunohistochemistry to assess GFP and mCherry expression in three different tumors derived from injection of a mixed population containing equal numbers of a primary and liver metastasis-derived pancreatic cancer cells into the liver as in (g), although the mixed population of cells were from an independent experiment where the primary tumor cells were labeled with mCherry and the liver metastasis cells were labeled with GFP. Scale bar, ~2500 μm. (**i**)

Immunohistochemistry to assess GFP, mCherry, or Cytokeratin-19 (Ck19) expression in whole mount liver tissue sections from a mouse from another independent experiment where a mixed population containing equal numbers of primary and liver metastasis-derived pancreatic cancer cells were implanted into the liver as in (g). (**j**) Representation of GFP-labeled primary or mCherry-labeled liver metastatic cells in a tumor derived from subcutaneous injection of mixed population containing equal numbers of labeled primary and liver metastasis-derived pancreatic cancer cells into the flank. Each data point represents one mouse; the numbers associated with each mouse indicate animal ID with each animal being injected with the same mixed population of cells. (**k**) Representation of mCherry-labeled primary or GFP-labeled liver metastatic cells in a tumor derived from subcutaneous injection of mixed population containing equal numbers of labeled primary and liver metastasis-derived pancreatic cancer cells into the flank. Each data point represents one mouse; the numbers associated with each mouse indicate animal ID with each animal being injected with the same mixed population of cells.

**Supplemental Figure 3**. **Assessment of glucose fate in primary pancreatic tumors and liver metastases*.*** (**a**) Plasma enrichment of U$^{13}$C-glucose as determined by GC-MS after a 6-hour U$^{13}$C-glucose infusion at a rate of 0.4 mg/min[51] in mice with a primary pancreatic tumor (pancreas; n=2 female mice) or with a liver metastatic pancreatic tumor (liver; n=4 female mice). (**b-h**) Fractional labeling of each indicated metabolite in the indicated tissue harvested from aged matched control mice (pancreas, liver) or from mice harboring a primary pancreatic tumor, a subcutaneous pancreatic tumor, or a liver metastatic tumor that was infused with U$^{13}$C-glucose as measured by GC-MS. In all cases tumors were generated by implanting pancreatic cancer cells derived from a primary tumor arising in the $KP^{-/-}C$ model. Mean +/- stdev; *p<0.5; **p<0.01; n.s.- not significant.

**Supplemental Figure 4. Assessment of glutamine fate in pancreatic tumors and liver metastases. (a-f)** Fractional labeling of each indicated metabolite in pancreatic cancer primary or liver metastasis tumors as measured by GC-MS following a 6-hour infusion of $U^{13}C$-glutamine at a rate of 3.7 mg/kg/min into mice. In all cases tumors were generated by implanting pancreatic cancer cells derived from a primary tumor arising in the $KP^{-/-}C$ mouse model. Mean +/- stdev; n=4 mice/group.

**Supplemental Figure 5. Amino acids levels in mouse pancreatic primary and liver metastatic tumors.** Levels of amino acids measured by LC-MS from primary pancreatic tumors and matched liver metastases arising in KPC mice; Mean +/- stdev (n=4).

**Supplemental Figure 6. Metabolic characterization of primary pancreatic cancer and liver metastatic cells in culture.** (**a**) Heatmap representation of unsupervised clustering of relative metabolite levels measured by LC-MS from paired primary and liver metastatic cells cultured in standard DMEM conditions; data is shown in triplicate per cell line from three independently derived cell lines. (**b-j**) Fractional labeling of each indicated metabolite in cells derived from a primary pancreatic tumor or a liver metastasis after being cultured with $U\text{-}^{13}C$ glucose for 24 hours. mean +/- stdev; data is shown from three independently derived cell lines; n=3.

**Supplemental Figure 7**. **Characterization of pancreatic tumor transplants into the pancreas, liver, and lung.** (**a**) Weights of pancreatic tumor tissue or liver tissue containing liver metastases (liver met) 4 weeks after implanting the same number of primary tumor- or liver metastasis-derived pancreatic cancer cells into the pancreas or the liver as indicated. Tissue weight +/- stdev; n=3-5 mice/condition. (**b**) Calculated tumor tissue weight (pancreas and liver) and tumor weight (subcutaneous) corresponding to data shown in Fig. 2b; tumor weight is calculated as the difference in tissue weight between the tumor-bearing organ and normal age-matched tissue from a non-tumor bearing mouse. For both pancreatic tumors and liver tumors, the normal tissue weight of the pancreas or the liver was subtracted to determine the calculated tumor weight, despite the observation that the entire pancreas was transformed with no macroscopic evidence of normal tissue. Subcutaneous (s.c.) tumors represent actual tumor weight. Mean+/- stdev; *$p < 0.05$, **$p < 0.01$; n=3-5 mice per condition. (**c**) Hematoxylin and Eosin (H&E) staining of tissue sections involving tumors arising from primary pancreas or liver metastatic cancer cells implanted in the pancreas or liver as indicated. The boundary between normal tissue and tumor is indicated. Scale bar- 3700 μm (lower magnification; 0.3x); 500 μm (higher magnification; 2x) (**d**) H&E staining and immunohistochemistry staining for CK19 in tumor tissue derived from primary or liver metastatic cancer cells that were implanted into the pancreas or the liver as indicated. scale bar, 50 μm. (**e**) Schematic depicting transplantation experiments to quantitatively assess pancreatic cancer cell proliferation in different tissue sites. (**f**) H&E staining of naturally arising KPC tumors (primary, liver and lung metastases (mets)) (top) or tumors resulting from transplantation of cells derived from cells isolated from the indicated primary, liver or lung metastatic tumors into each indicated site (bottom); scale bar, 50 μm.

**Supplemental Figure 8. Assessment of proliferation and cell death in tumors.** (**a-c**) Ki67 staining of tumor tissue derived from primary or liver metastatic (Liver Met) pancreatic cancer cells that were implanted into the pancreas (left) or the liver (right) (a) or staining of tumor tissue from primary or lung metastatic (Lung Met) pancreatic cancer cells implanted into the pancreas (b) or lung (c) as indicated. scale bar, 50 μm. (**d**) Immunohistochemistry stain for BrdU in a representative tumor obtained from primary or liver metastatic pancreatic cancer cells injected in either the pancreas (left) or liver (right). scale bar, 20 μm. (**e-g**) Immunohistochemistry staining for cleaved caspase 3 in tumor sections derived from primary, liver, or lung metastatic pancreatic cancer cells injected in either the pancreas (e), liver (f) or lung (g). Representative region per section is shown with higher magnification image shown in the bottom right inset; scale bar for panels e-g: lower magnification- 200 μm, inset- 20 μm.

**Supplemental Figure 9. Independent assessment of tumor growth in pancreas and liver in immunocompetent and nude mice.** (**a**) Weights of the indicated tissues harvested from mice where primary, or liver metastatic pancreatic cancer cells were implanted into the pancreas, liver, or subcutaneous space on the flank. Weight of age-matched normal pancreatic and liver tissue is also shown. n=3-5 male and female mice; Mean+/- stdev; n.s.- not significant, *p <0.05. Also shown (right) is the calculated tumor weight (pancreas and liver) and tumor weight (subcutaneous; s.c.) for these data with tumor weight calculated as the difference in tissue weight between the tumor-bearing organ and the normal age-matched tissue from a non-tumor bearing mouse. For both pancreatic tumors and liver tumors, the normal tissue weight of the pancreas or the liver was subtracted to determine the calculated tumor weight, despite the observation that the entire pancreas was transformed with no macroscopic evidence of normal tissue. Subcutaneous (s.c.) tumors represent actual tumor weight. mean+/- stdev; ***p< 0.001. (**b**) Calculated tumor weight represented as the difference in tissue weight between tumor-bearing organ and normal age-matched tissue from non-tumor bearing age-matched nude (NU/J) mice; (n=3-4 mice/group) (**c**) Tissue weight from Fig. 2h, normalized to mouse body weight. (n=3-4 mice/group); ***p<0.001, *p<0.05, n.s.- not significant; mean+/- stdev (left); calculated tumor weight (as in (a)) normalized to body weight (n=3-4 mice/group); *p<0.05, n.s.- not significant; mean+/- stdev (right). (**d**) Weight of mice 4 weeks post-implantation of the indicated cancer cells (primary or liver met; tumor bearing mice) or sham injection (non-tumor bearing mice) into the pancreas (left) or liver (right) as indicated; n=3-4 mice/group; mean+/- stdev.

**Supplemental Figure 10. Characterization of pancreatic tumor transplants into the pancreas, liver, and lung.** (**a**) Weights of the indicated tissues harvested from mice where primary pancreatic cancer cells, or liver or lung metastatic pancreatic cancer cells were implanted into the pancreas (pancreatic tumor), liver (liver+tumor), or lung (lung+tumor). Tissue weight +/- stdev. n=3-5 mice. (**b**) Calculated tumor weight (pancreas, liver, and lung) and tumor weight (subcutaneous) corresponding to data shown in Fig. 2k; tumor weight is calculated as the difference in tissue weight between the tumor-bearing organ and normal age-matched tissue from a non-tumor bearing mouse. For pancreatic tumors, liver tumors, and lung tumors the normal tissue weight of the pancreas, the liver, or the lung was subtracted to determine the calculated tumor weight, despite the observation that the entire pancreas was transformed with no macroscopic evidence of normal tissue. Subcutaneous tumors (s.c.) represent actual tumor weight. mean+/- stdev; *p<0.05, *p<0.01, ***p< 0.001. n=3-5 mice. (**c**) Macroscopic images of liver (left) or lung (right) four weeks post injection of cancer cells derived from a primary pancreatic tumor (primary >) or from a liver or lung metastasis (liver met > or lung met >, respectively) into the indicated organ. N- normal tissue; T- tumor; arrows indicate tumor nodules in the lung. (**d-f**) Whole mount H&E-stained liver or lung tissue from mice where primary pancreatic cancer cells (d), liver (e), or lung (f) metastatic cancer cells were implanted into the indicated site. Scale bars are indicated in the figure panels; higher magnification images are shown for smaller tumors; tumors in the liver are indicated.

**Supplemental Figure 11. Passaging pancreatic tumors in the lung or liver of mice selects for increased seeding of the metastatic tissue.** (**a**) Macroscopic tissue images and histology assessing that tissue after each passage where cells from a lung metastasis (top) or liver metastasis (bottom) were serially implanted to from tumors in each organ. P1-P3 refers to each passage. Tumor areas indicated by boundary or arrows. (**b**) Quantification of lung metastases generated when parental (P0) or P3 lung metastatic pancreatic cancer cells were injected into the tail vein of wild type mice (left). Tumor area was calculated as the area of the tumor divided by the total area of tissue for all lung sections as assessed by H&E staining. Mean +/- stdev; n=3 mice/condition. (**c**) Quantification of the number of lung nodules generated when parental (P0) or P3 lung metastatic pancreatic cancer cells were implanted to form a tumor in the pancreas (right). Mean +/- stdev; n=3-4 mice/condition. (**d**) Quantification of liver metastases generated when parental (P0) or P3 liver metastatic pancreatic cancer cells were injected into the liver. Tumor area was calculated as the area of the tumor divided by the total area of tissue for all liver sections as assessed by H&E staining; Mean +/- stdev; n=3-4 mice/condition. (**e**) Macroscopic images of liver (top) or lung and liver (bottom) after implantation of P3 liver metastatic pancreatic cancer cells (Liver Met P3 >) into the Liver (top) or the tail vein (bottom) as indicated. N- normal tissue; T- tumor; arrows indicate tumor nodules in the lung or liver. (**f**) Calculated tumor tissue weight (pancreas, liver, and lung) and tumor weight (subcutaneous) corresponding to data shown in Fig. 3b-e; tumor weight is calculated as the difference in tissue weight between the tumor-bearing organ and normal age-matched tissue from a non-tumor bearing mouse. Subcutaneous tumors represent actual tumor weight. For pancreatic tumors, liver tumors, and lung tumors the normal tissue weight of the pancreas, the liver, or the lung was subtracted to determine the calculated tumor weight, despite the observation that the entire pancreas was transformed with no macroscopic evidence of normal tissue. Mean+/- stdev; n=3-5 mice/condition *$p<0.05$, **$p<0.01$, ***$p<0.001$; n.s.- not significant

**Supplemental Figure 12. Analysis of tumors derived from primary liver and primary lung cancer cells.** (**a**) Calculated tumor weight corresponding to relevant data shown in Fig. 3b-d and Fig. 4a. Tumor weight is calculated as the difference in tissue weight between the tumor-bearing organ and normal age-matched tissue from a non-tumor bearing mouse. For pancreatic tumors, liver tumors, and lung tumors the normal tissue weight of the pancreas, the liver, or the lung was subtracted to determine the calculated tumor weight. Mean+/- stdev; n=3-4 mice/condition *p<0.05, **p<0.01, ***p<0.001; n.s.- not significant. (**b**) Histology of tumor sections corresponding to intratracheal injection of indicated cancer cells into the lung that relate to data shown in Fig 4b. (**c**) Ultrasound guided injection (USGI) of primary lung cancer cells or pancreatic cancer lung metastases into the lung and quantification of tumor area. (**d**) Representative histology of tumor sections that related to data shown in (c) showing single tumor nodule formation in the lung following UGSI. (**e**) Calculated tumor weight corresponding to data shown in Fig. 4c as well as data in fig. 3b-d. Tumor weight is calculated as the difference in tissue weight between the tumor-bearing organ and normal age-matched tissue from a non-tumor bearing mouse. For pancreatic tumors and liver tumors the normal tissue weight of the pancreas and the liver was subtracted to determine the calculated tumor weight. Mean+/- stdev; n=3-4 mice/condition *p<0.05.

**Supplemental Figure 13. Histology images of primary lung and primary liver tumors.** (**a-b**) H&E stained histology sections of tumors obtained from injection of primary lung cancer cells into either the lung (top) or liver (bottom) (a) or from injection of primary liver cancer cells (HCC) into the lung (top) or liver (bottom) (b). n=3 mice/condition; data accompany tumor area data shown in Fig. 4d-e.

**Supplemental Figure 14. Assessment of metabolite levels in tissues.** (**a**) Principal component analysis (PCA) plot showing the separation of metabolite levels measured in pooled in interstitial fluid form indicated organs or plasma from wild-type non-tumor bearing animals. (**b**) Proliferation rate of primary lung cancer cells (human A549 cells or mouse KP primary lung cancer cells) and cells from a KPC pancreatic cancer lung metastasis (lung met) in media formulated to mimic metabolite levels measured in lung interstitial fluid (lung media).

**Supplemental Figure 15. Analysis of metabolic gene expression from mouse and human tissues.** (**a-c**) Metabolic gene expression derived from scRNA-seq data[54] that is another representation of the data presented in figure 4g-h. Data represented as entire cell populations and comparing primary pancreatic cancer (PDAC) and liver metastases (liver met) arising in the mouse KPC PDAC model (a); A cross correlation of metabolic gene expression between pairs of single cells are plotted ordered by either site (b), or barcode (c); within a given barcode, pairs of cells tend to have high correlations in their metabolic expression, whereas there are subsets of cells within sites that have different metabolic gene expression but these subsets of cells do not separate based on tissue site; i.e. a given barcode across has similar metabolic gene expression across metastatic sites. (**d**) PCA of a human RNA-seq dataset[6,7] comparing metabolic gene expression in primary pancreatic tumors (Primary PDAC; green), liver metastatic PDAC (yellow), hepatocellular carcinoma (HCC; purple), and healthy liver (blue) as indicated. (**e**) Average Pearson correlation comparing metabolic similarity between pancreatic liver metastases (Met PDAC; yellow data points) to primary tumors (Primary PDAC), hepatocellular carcinoma

(HCC) or healthy liver. Liver metastatic PDAC with higher Pearson correlation to HCC and healthy liver contains a higher degree of hepatocyte contamination (circled yellow data points) in (d) and (e).
