## Supplemental Figures for "Cancer tissue of origin constrains the growth and metabolism of metastases"

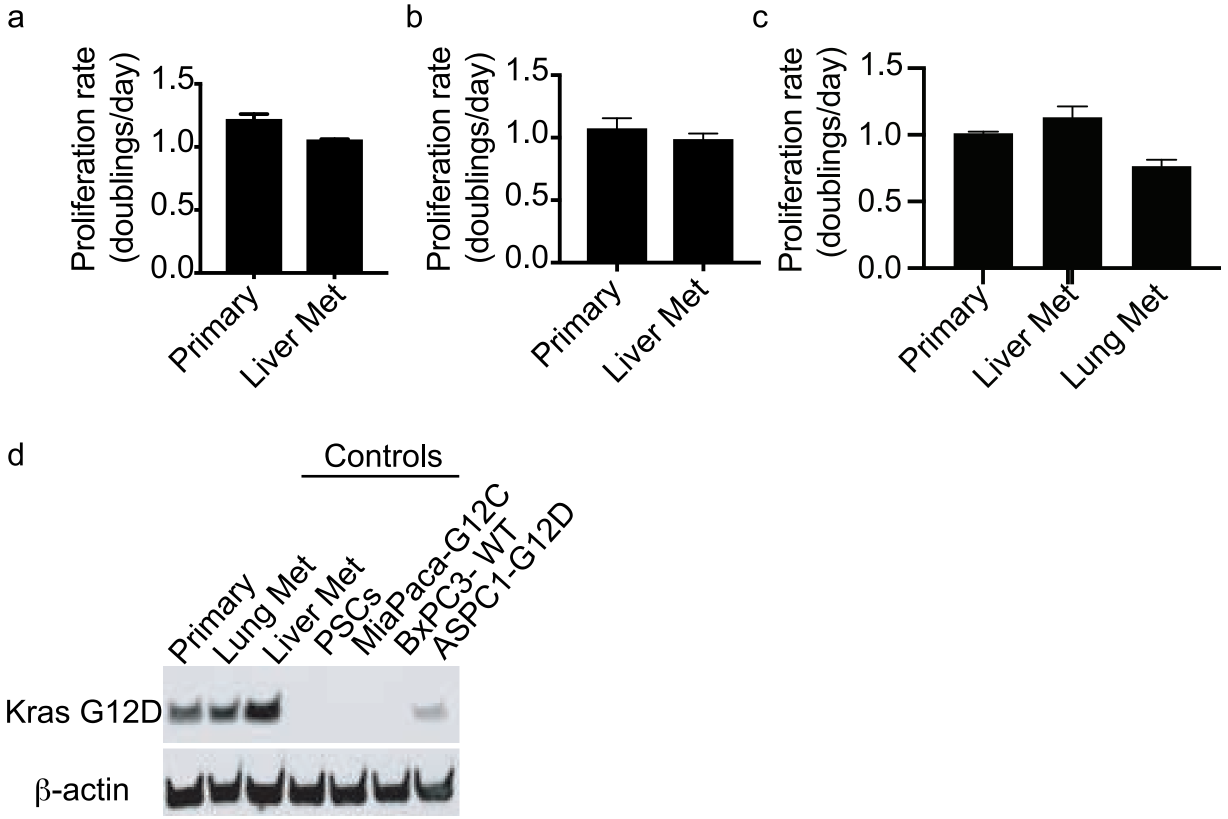
a
b
c
d
Proliferation rate (doublings/day)
1.5
1.0
0.5
0.0
Primary
Liver Met
Lung Met
Controls
Primary
Lung Met
Liver Met
PSCs
MiaPaca-G12C
BxPC3- WT
ASPC1-G12D
Kras G12D
β-actin

Supplemental Figure 2

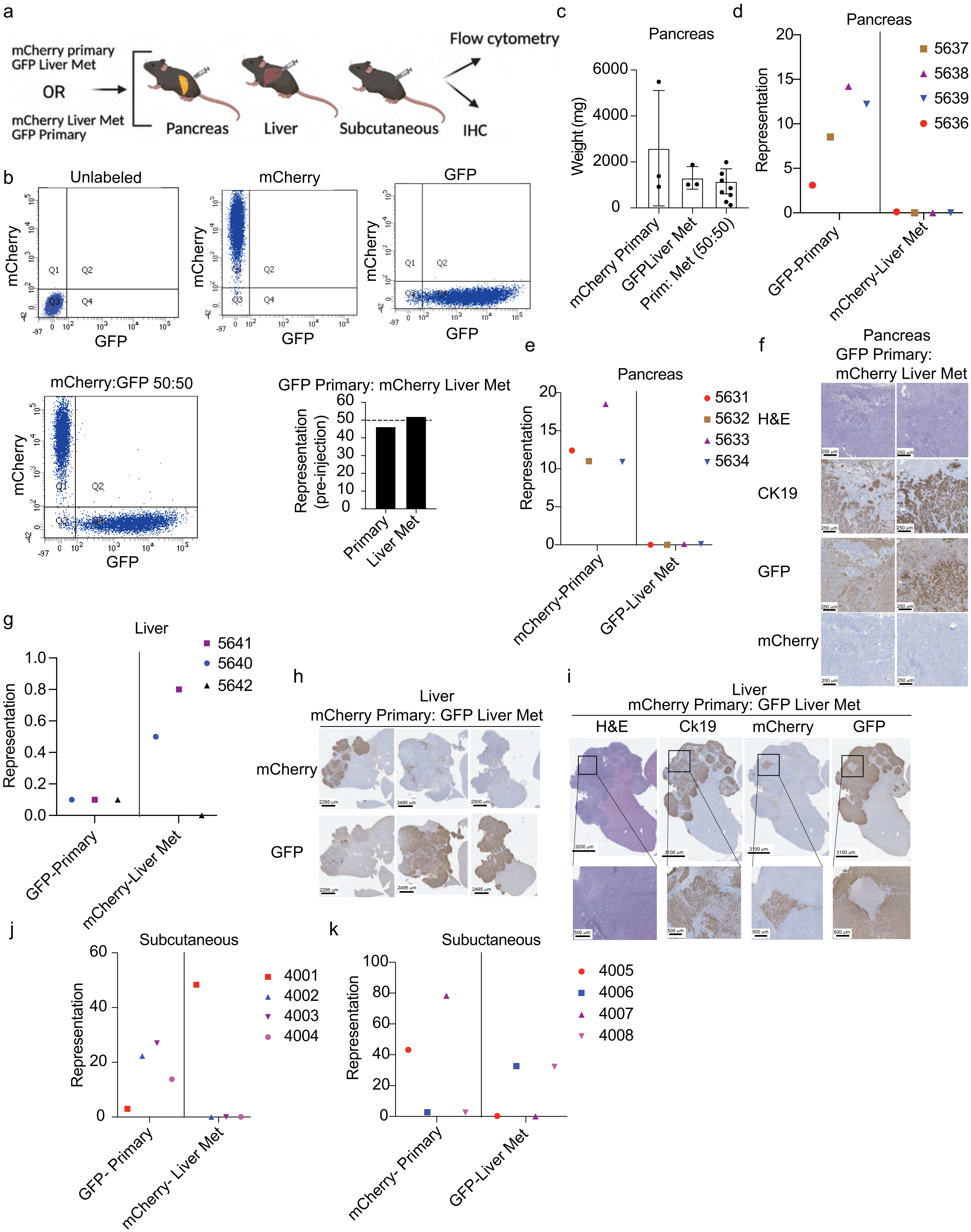

Supplemental Figure 3

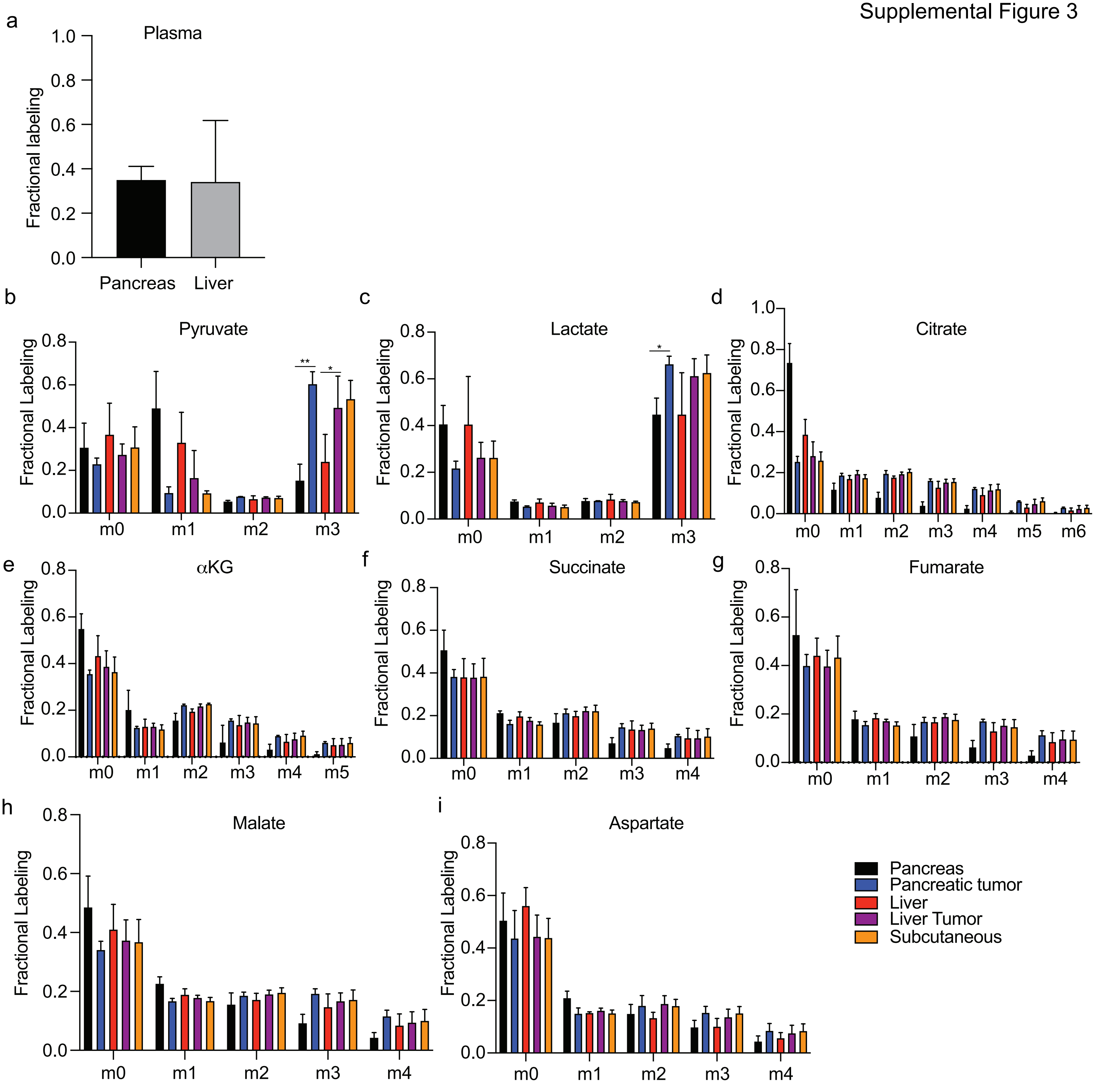

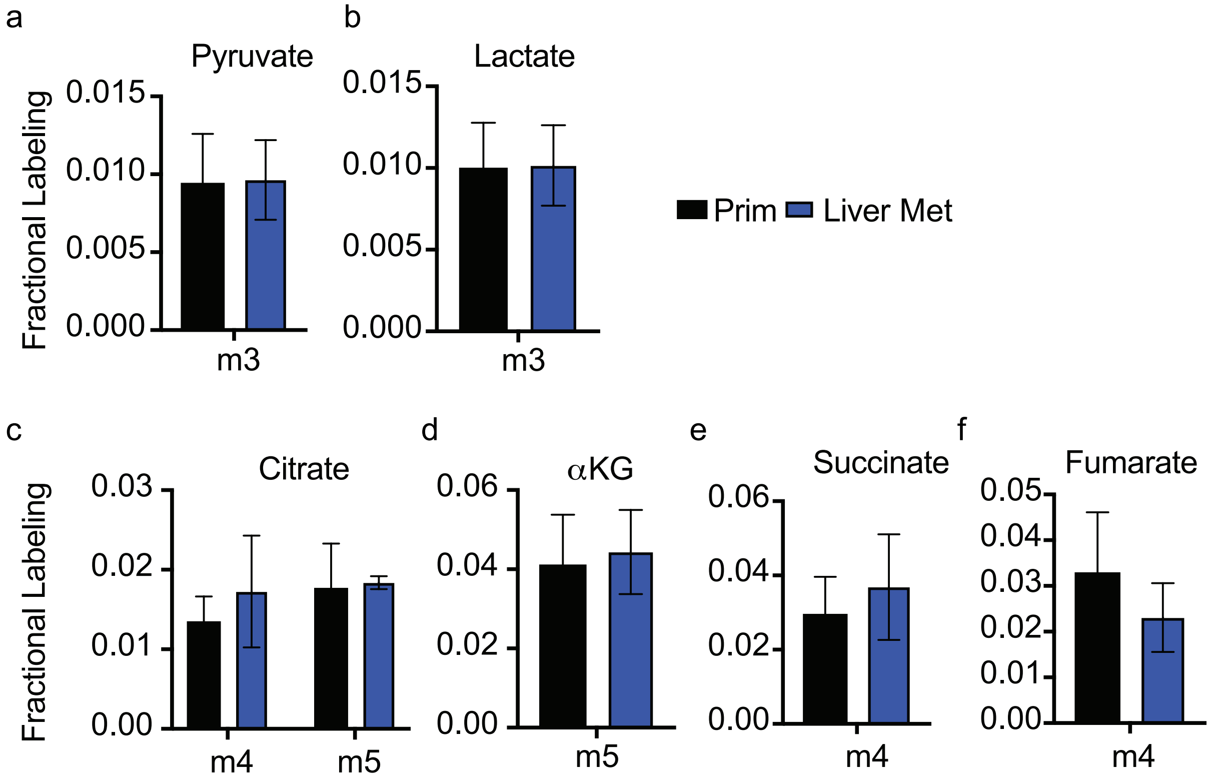
a
Pyruvate
Fractional Labeling
0.015
0.010
0.005
0.000
m3
b
Lactate
0.015
0.010
0.005
0.000
m3
Prim
Liver Met
c
Citrate
Fractional Labeling
0.03
0.02
0.01
0.00
m4
m5
d
αKG
0.06
0.04
0.02
0.00
m5
e
Succinate
0.06
0.04
0.02
0.00
m4
f
Fumarate
0.05
0.04
0.03
0.02
0.01
0.00
m4

Supplemental Figure 5

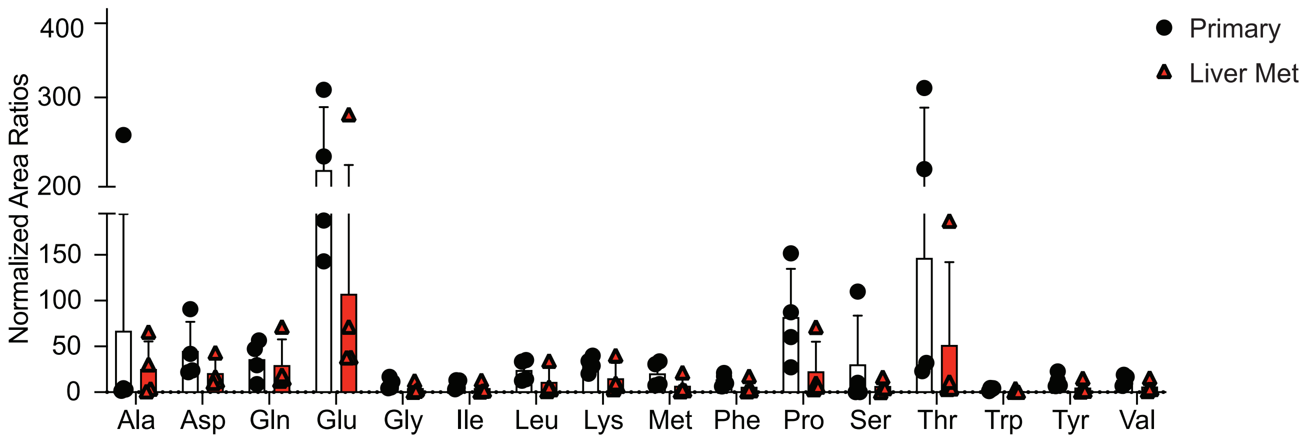

Supplemental Figure 6

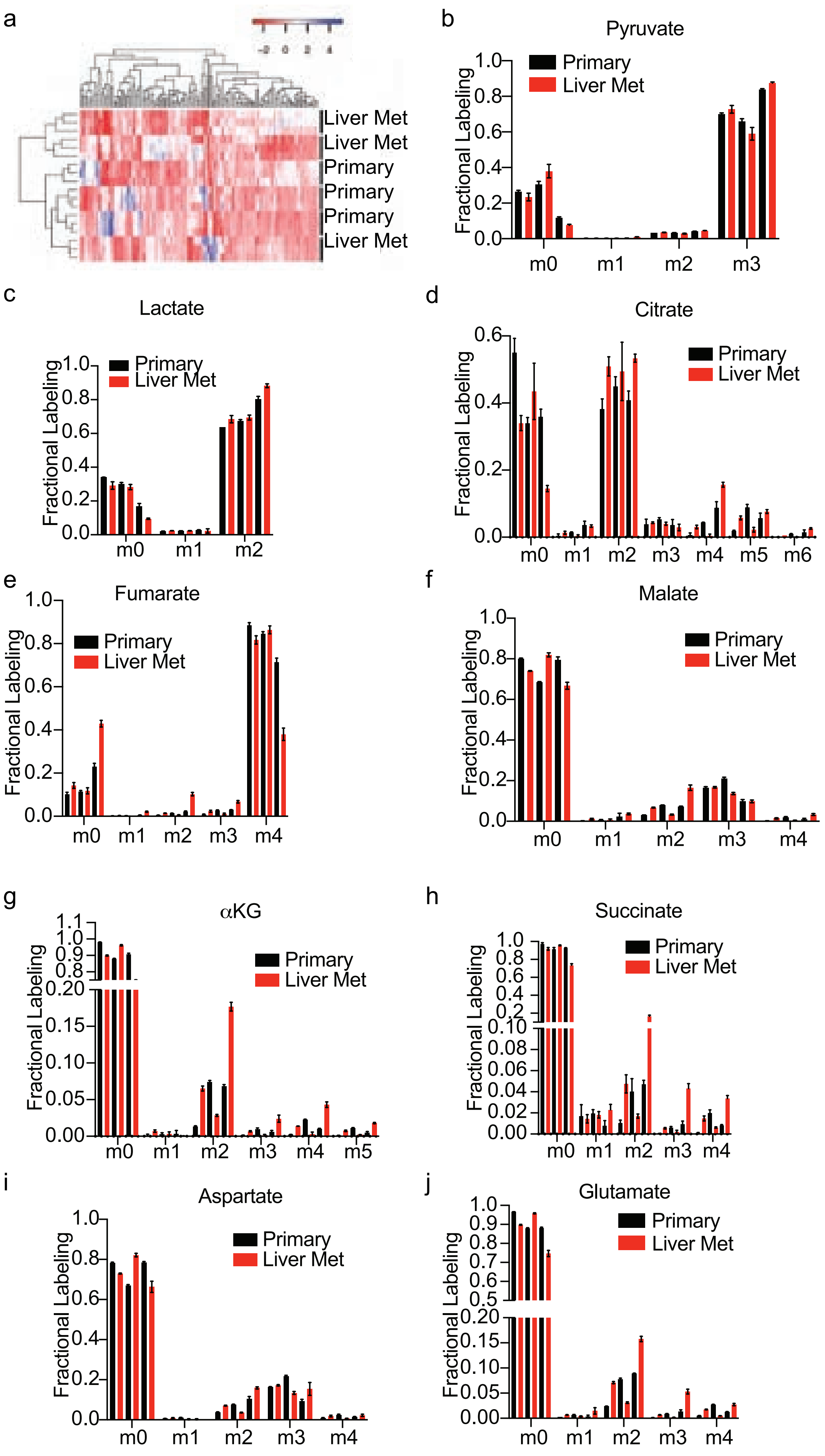

a

| | Pancreatic Tumor (mg) | Liver + Tumor (mg) |
|---|---|---|
| Primary | 1347 ± 361 | 1444 ± 518 |
| Liver Met | 1987 ± 400 | 1376 ± 292 |

b

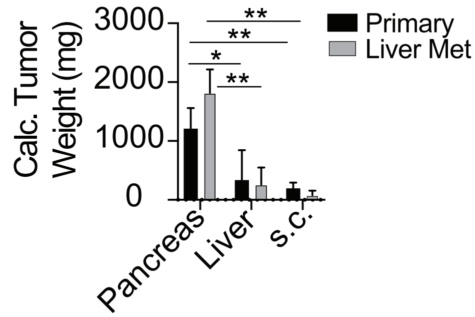


c

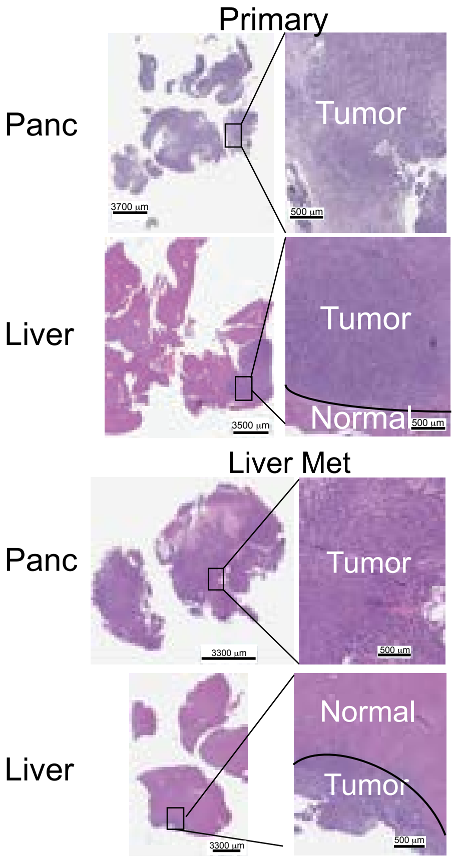


d

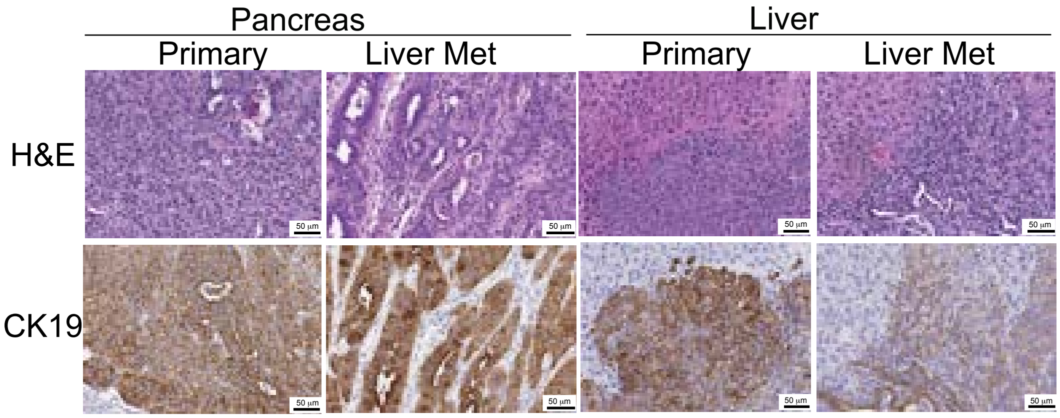


e

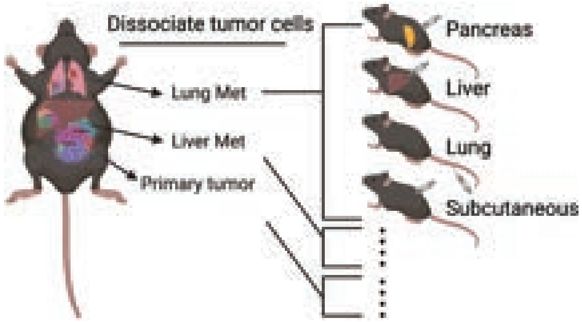


f

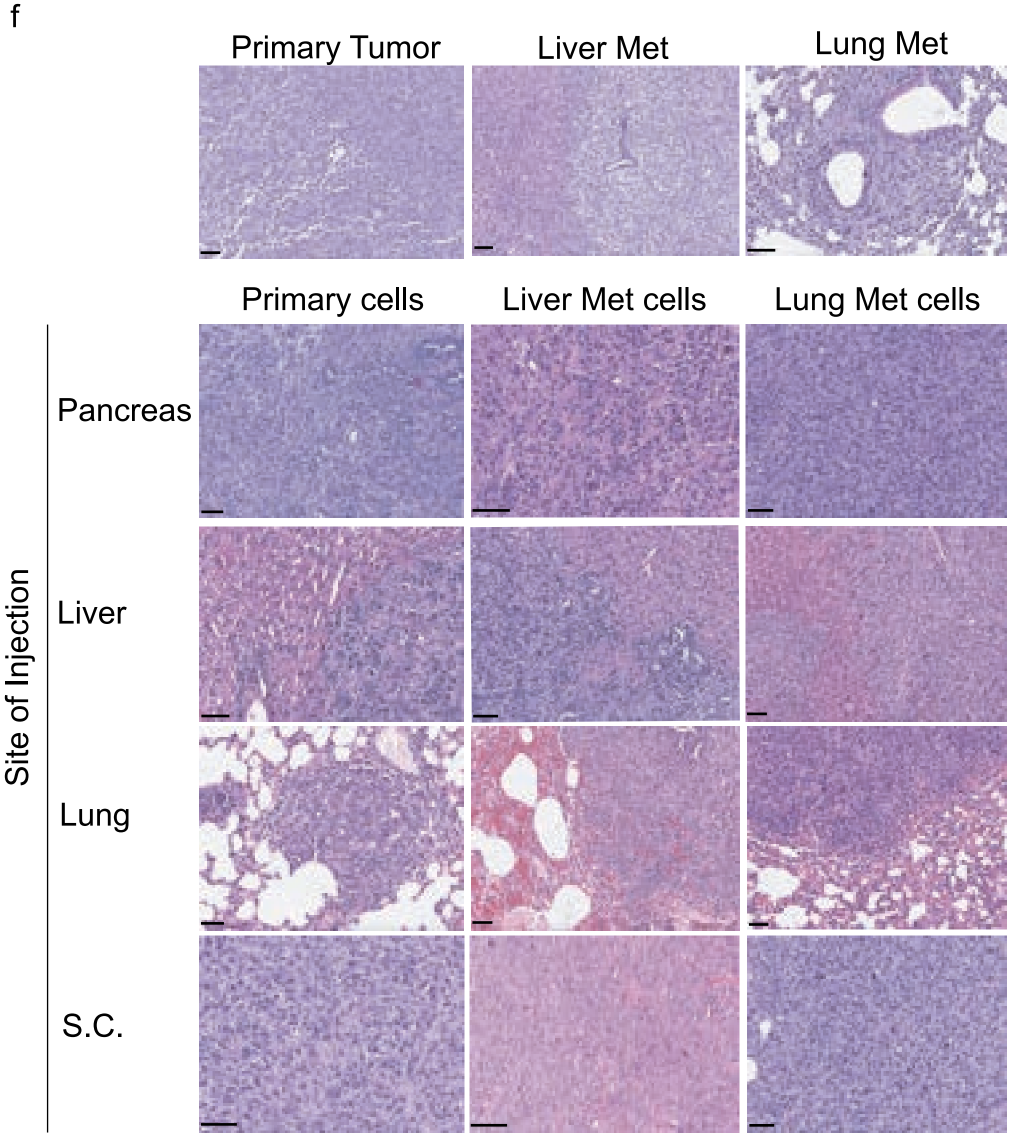

Supplemental Figure 8

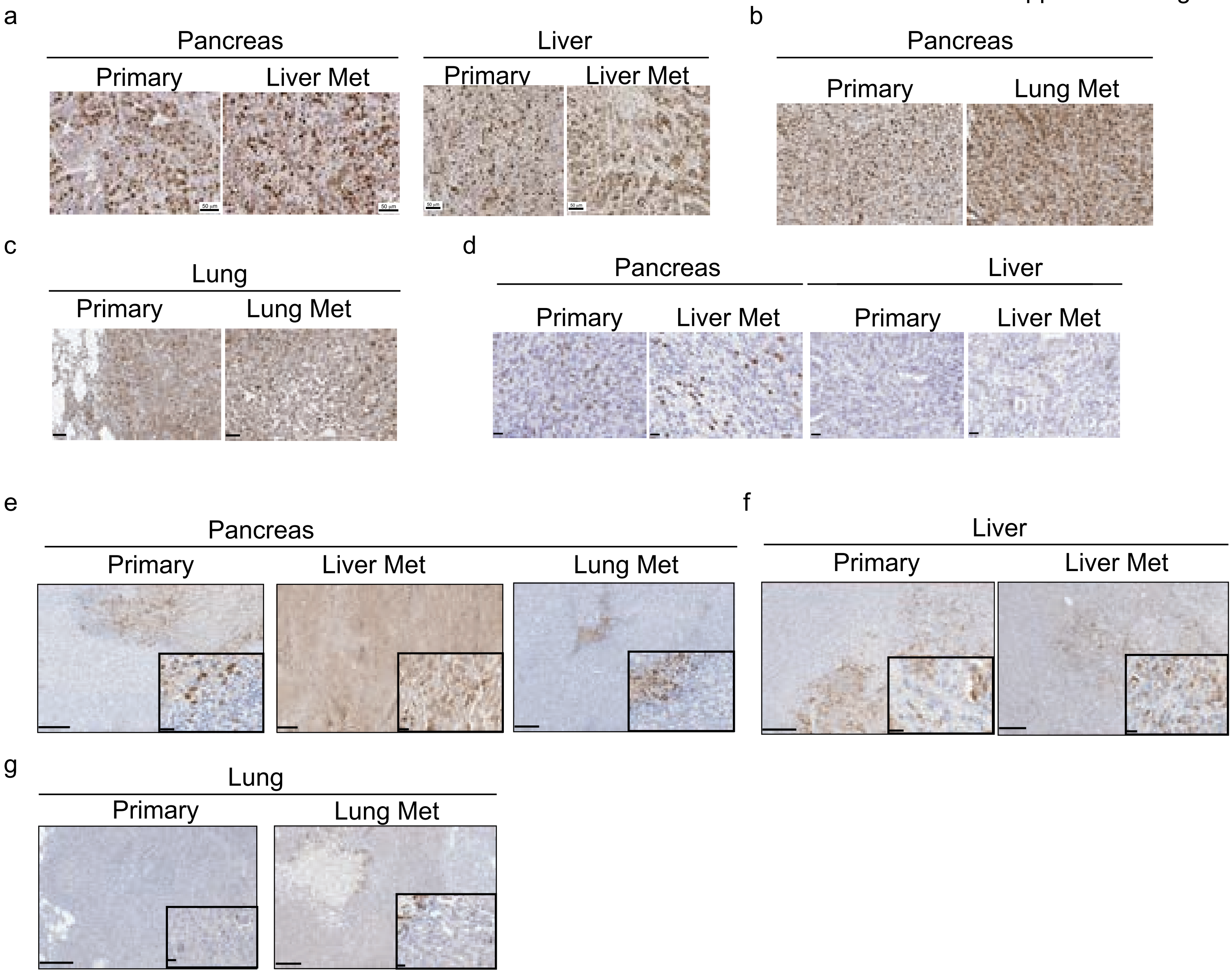

Supplemental Figure 9

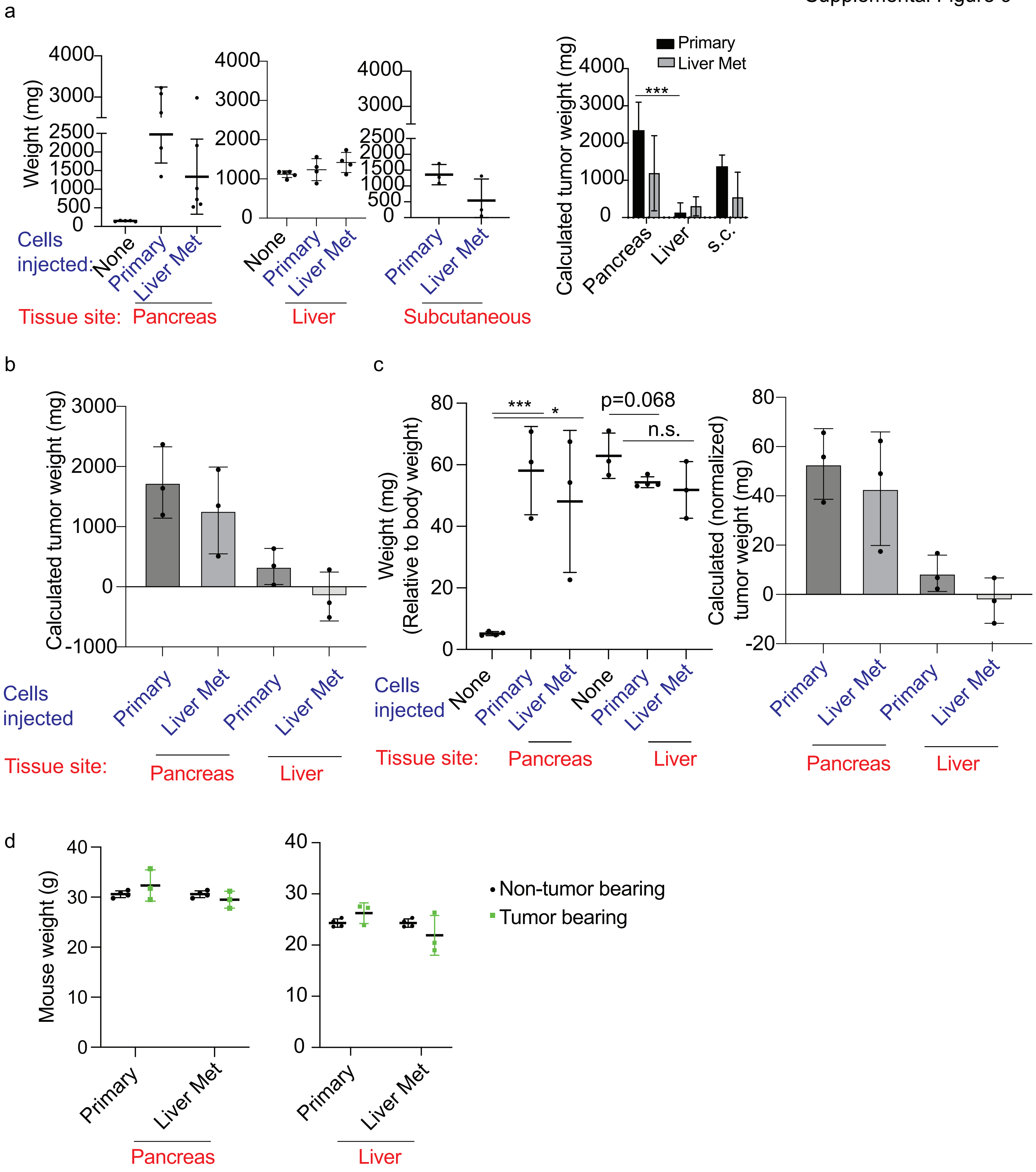

Supplemental Figure 10

a

| | Weight (mg) | | |
|---|---|---|---|
| | Pancreatic Tumor | Liver + Tumor | Lung + Tumor |
| Primary | 1376 ± 680 | 1408 ± 212 | 360 ± 180 |
| Liver Met | 1158 ± 366 | 1442 ± 270 | n/a |
| Lung Met | 764 ± 212 | n/a | 272 ± 48 |

b

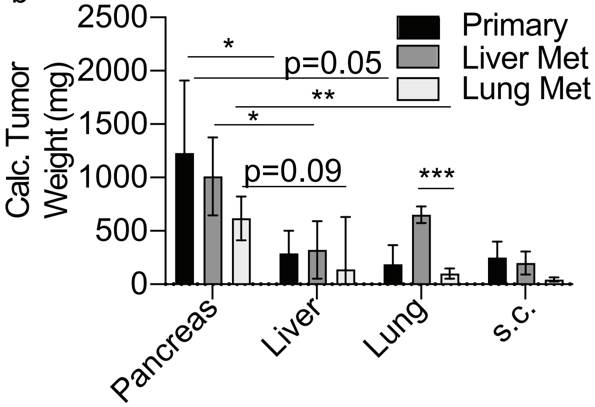


c

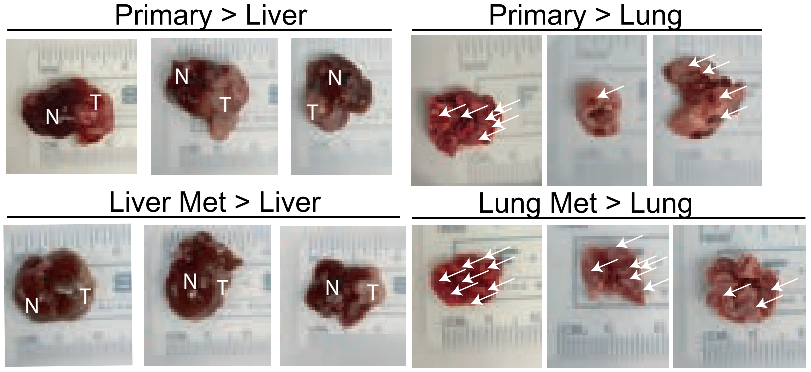


d

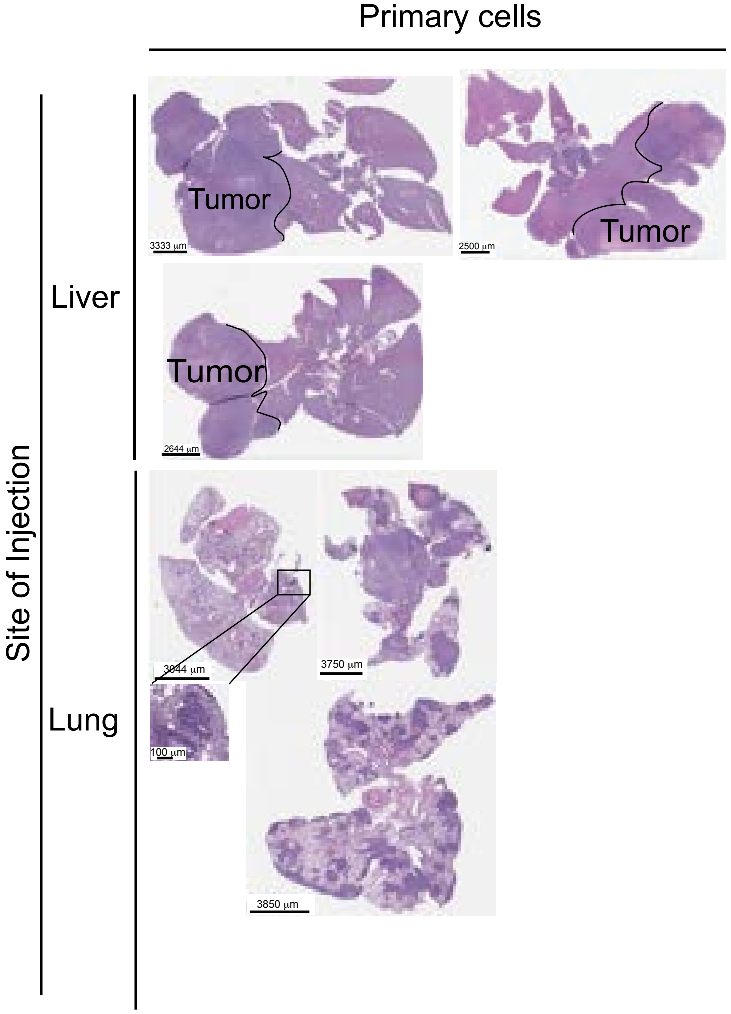


e

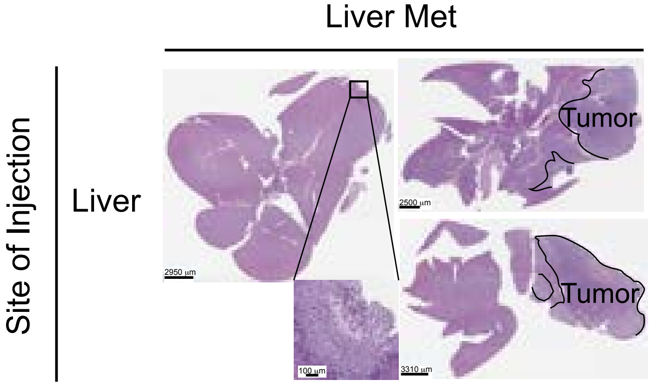


f

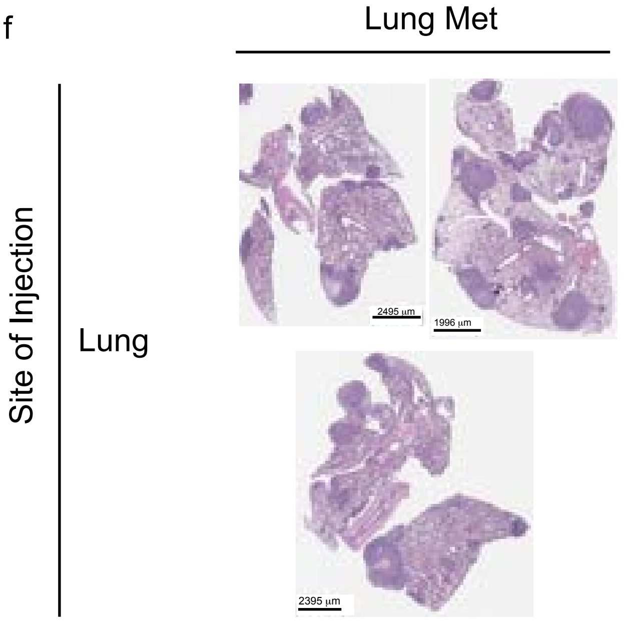

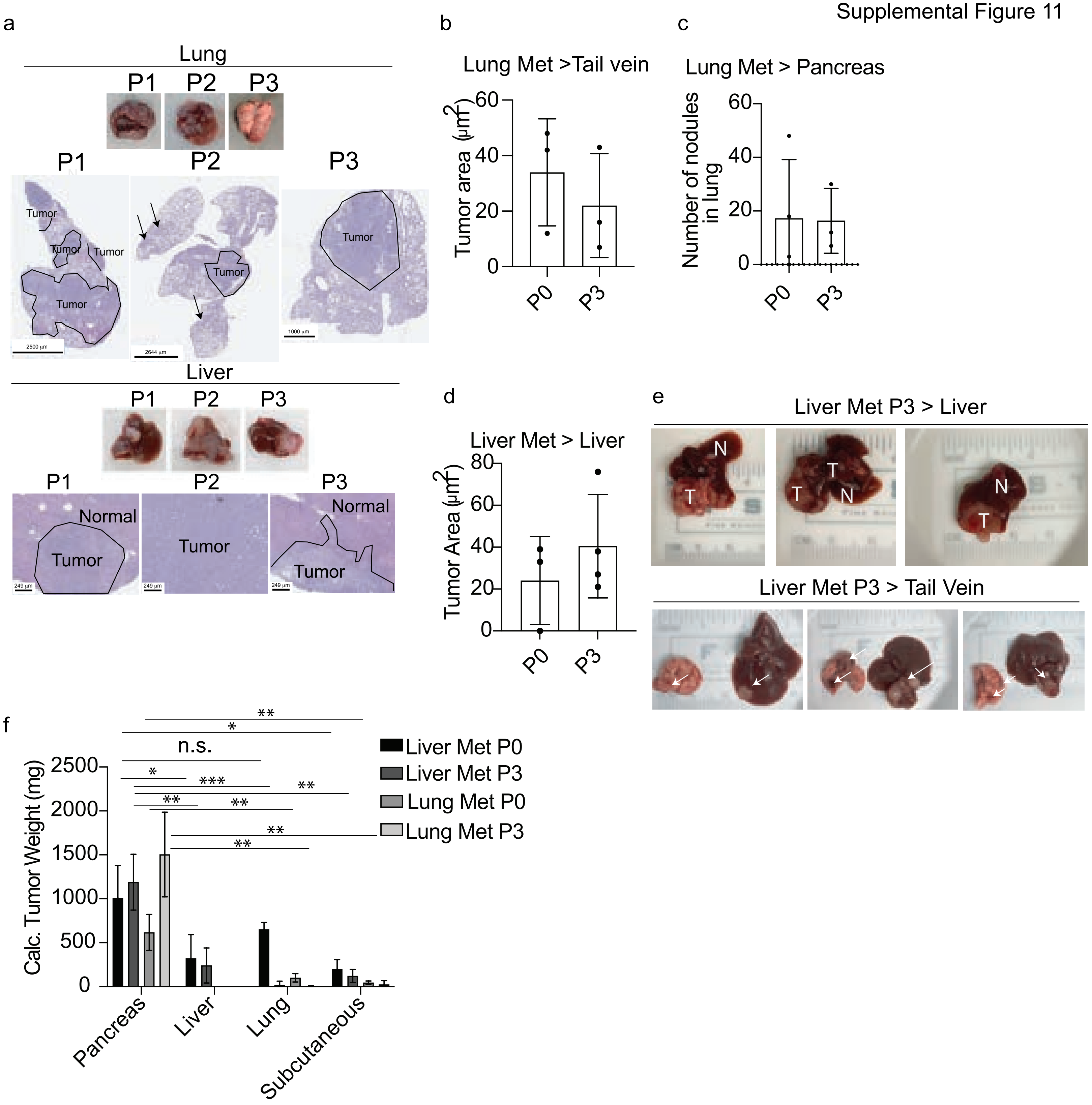
a
Lung
P1
P2
P3
Tumor
Liver
Normal
b
Lung Met >Tail vein
Tumor area (μm2)
P0
P3
c
Lung Met > Pancreas
Number of nodules in lung
d
Liver Met > Liver
Tumor Area (μm2)
e
Liver Met P3 > Liver
T
N
Liver Met P3 > Tail Vein
f
Calc. Tumor Weight (mg)
Liver Met P0
Liver Met P3
Lung Met P0
Lung Met P3
n.s.
Pancreas
Liver
Lung
Subcutaneous

Supplemental Figure 12

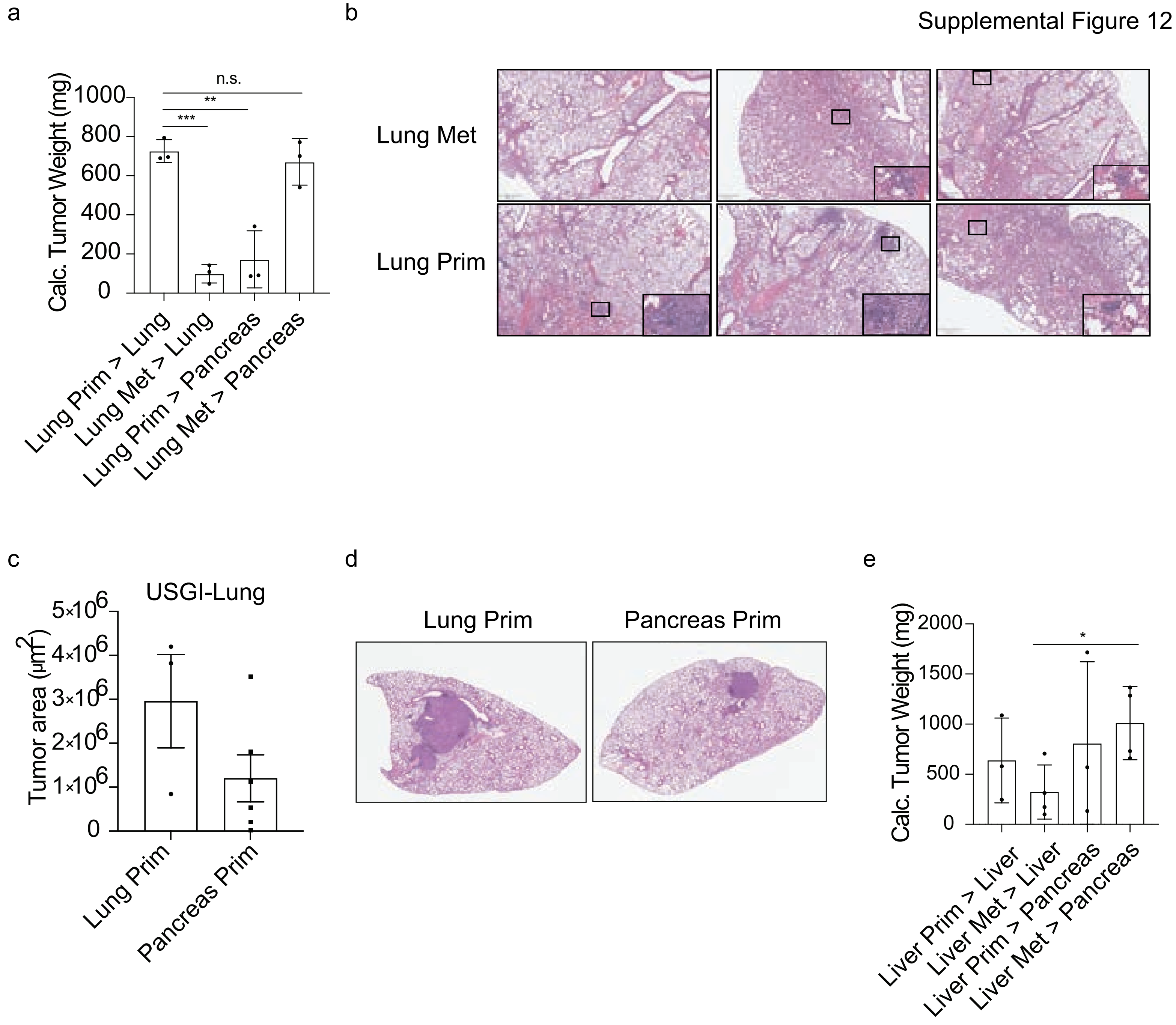

a

Lung adenocarcinoma

Lung

Liver

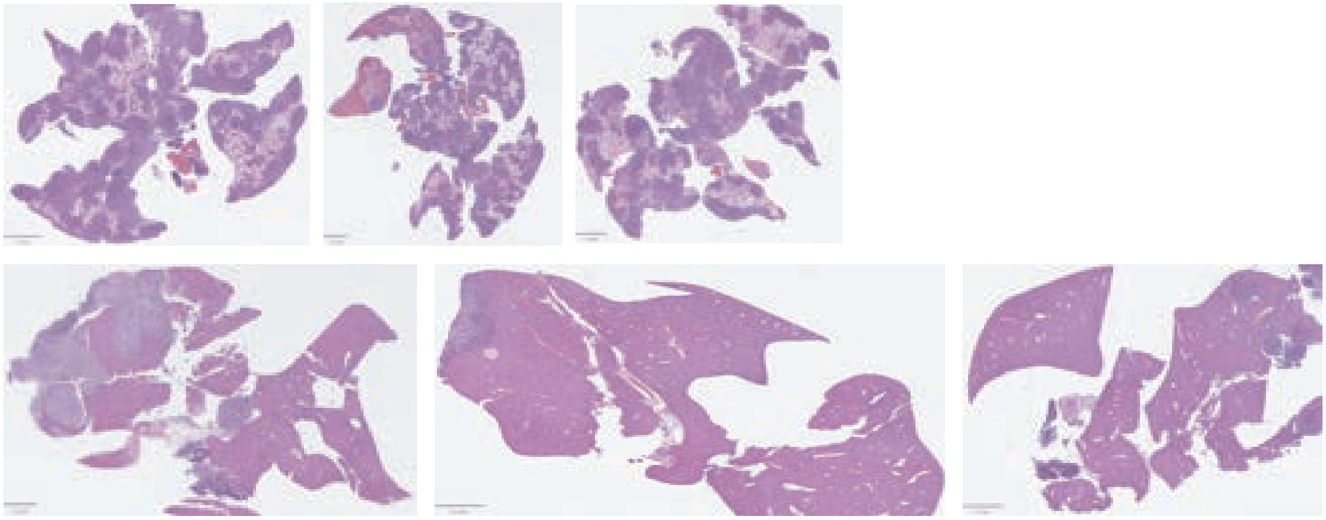

b

HCC

Lung

Liver

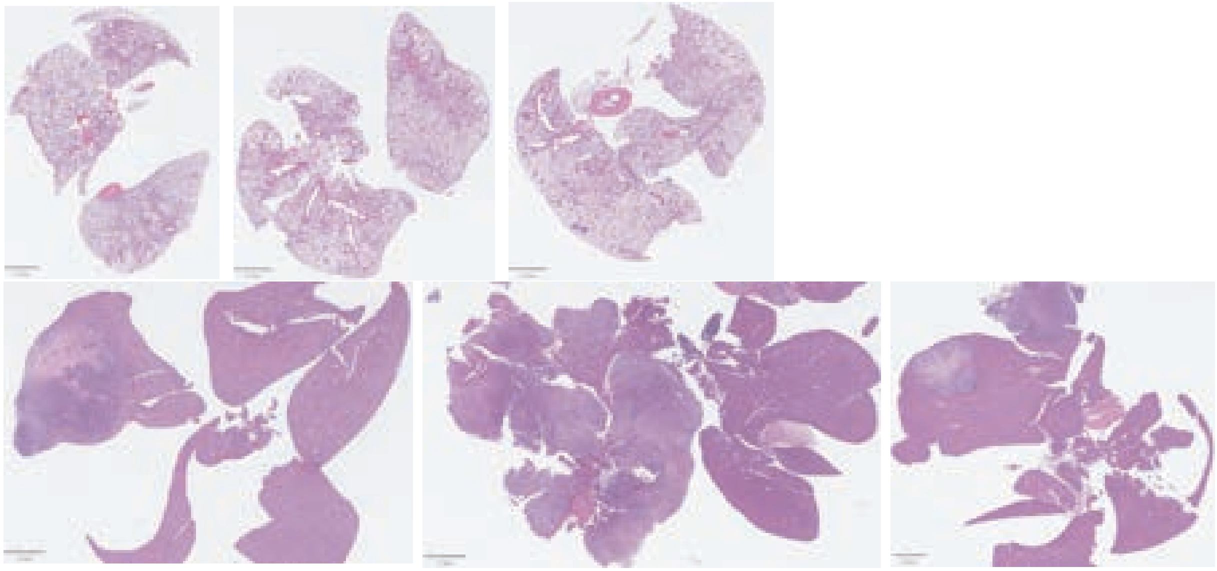

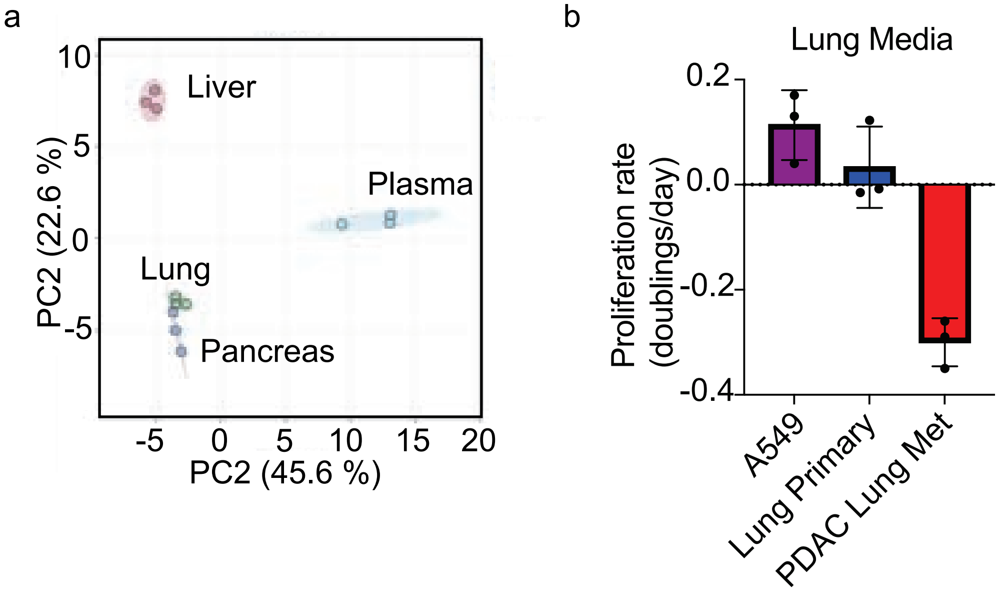
a
Liver
Plasma
Lung
Pancreas
PC2 (22.6 %)
PC2 (45.6 %)
10
5
0
-5
-5
0
5
10
15
20
b
Lung Media
Proliferation rate (doublings/day)
0.2
0.0
-0.2
-0.4
A549
Lung Primary
PDAC Lung Met

Supplemental Figure 15

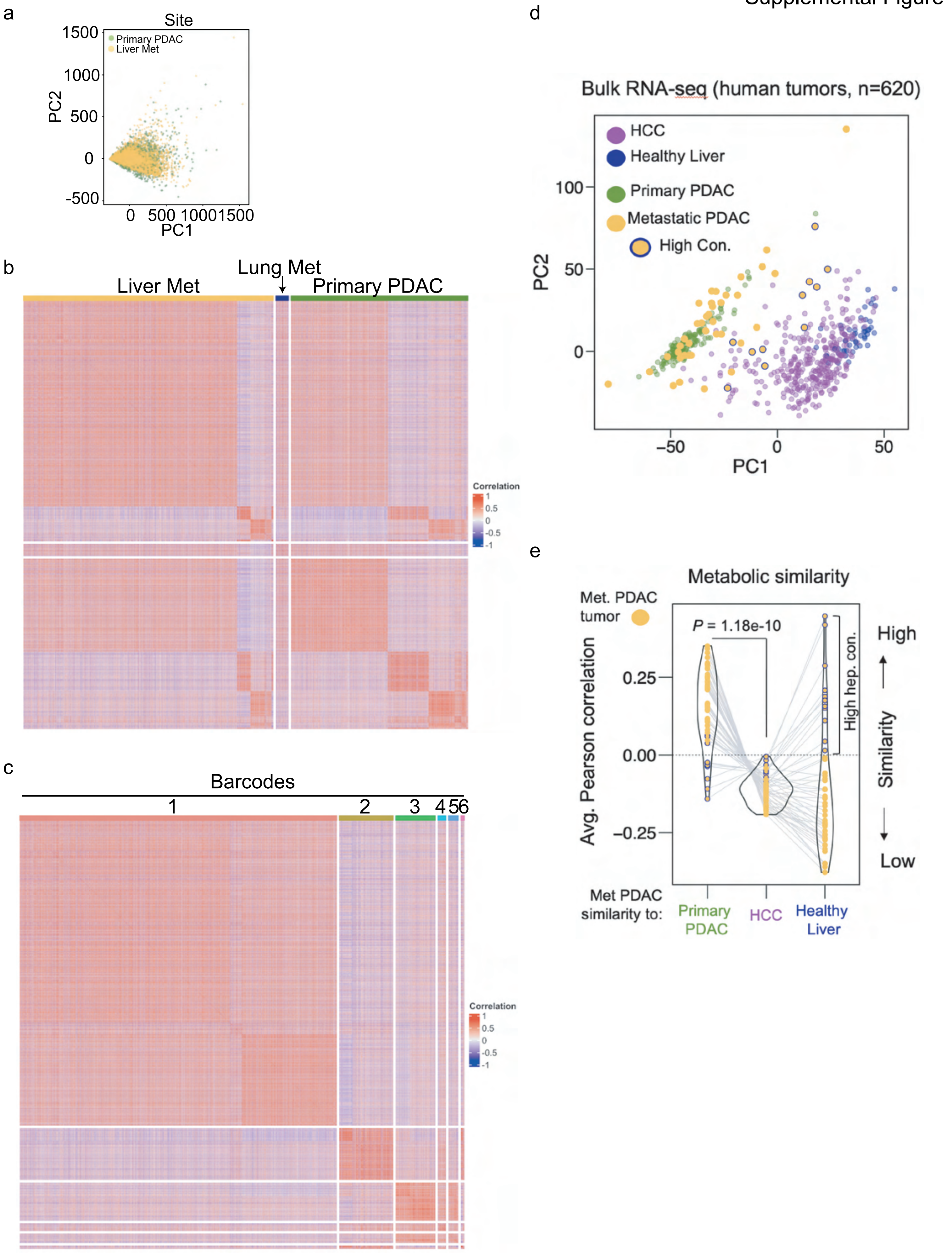
